# The dynamic effect of regulatory genetic variation on the *in vivo* ER stress transcriptional response

**DOI:** 10.1101/2021.03.12.435172

**Authors:** Nikki D. Russell, Clement Y. Chow

## Abstract

Genotype x Environment (GxE) interactions occur when environmental conditions drastically change the effect of a genetic variant. In order to truly understand the effect of genetic variation, we need to incorporate multiple environments into our analyses. Many variants, under steady state conditions, may be silent or even have the opposite effect under stress conditions. This study uses an *in vivo* mouse model to investigate how the effect of genetic variation changes with tissue type and cellular stress. Endoplasmic reticulum (ER) stress occurs when misfolded proteins accumulate in the ER. This triggers the unfolded protein response (UPR), a large transcriptional response which attempts to return the cell to homeostasis. This transcriptional response, despite being a well conserved, basic cellular process, is highly variable across different genetic backgrounds, making it an ideal system to study GxE effects. In this study, we sought to better understand how genetic variation alters expression across tissues, in the presence and absence of ER stress. The use of different mouse strains and their F1s allow us to also identify context specific *cis*- and *trans*-regulatory mechanisms underlying variable transcriptional responses. We found hundreds of genes that respond to ER stress in a tissue- and/or genotype-dependent manner. Genotype-dependent ER stress-responsive genes are enriched for processes such as protein folding, apoptosis, and protein transport, indicating that some of the variability occurs in canonical ER stress factors. The majority of regulatory mechanisms underlying these variable transcriptional responses derive from *cis*-regulatory variation and are unique to a given tissue or ER stress state. This study demonstrates the need for incorporating multiple environments in future studies to better elucidate the effect of any particular genetic factor in basic biological pathways, like the ER stress response.

**Author Summary:** The effect of genetic variation is dependent on environmental context. Here we use genetically diverse mouse strains to understand how genetic variation interacts with stress state to produce variable transcriptional profiles. In this study, we take advantage of the endoplasmic reticulum (ER) stress response which is a large transcriptional response to misfolded proteins. Using this system, we uncovered tissue- and ER stress-specific effects of genetic variation on gene expression. Genes with genotype-dependent variable expression levels in response to ER stress were enriched for canonical ER stress functions, such as protein folding and transport. These variable effects of genetic variation are driven by unique sets of regulatory variation that are only active under context-specific circumstances. The results of this study highlight the importance of including multiple environments and genetic backgrounds when studying the ER stress response and other cellular pathways.

## Introduction

Genetic variation rarely acts in isolation and changes in environmental conditions can drastically alter the effect of a particular variant. This genotype x environment (GxE) effect is pervasive in biology, where the environment can be broadly defined as anything that is not genetic, including external conditions, tissue type, and cellular stressors. A GxE effect is defined as an interaction between multiple genotypes and environmental factors, where genotypes respond to different environments in a non-additive way. GxE effects are particularly evident in RNA levels where context-specific changes in expression are the norm. GxE studies are most commonly performed with genetically diverse individuals and a single change in context. For example, the GTEx consortium has made significant progress in understanding the genetic architecture underlying gene expression across tissues in healthy human individuals [1]. It is rare, however, for GxE studies to incorporate multiple contexts, including environment, tissue type, and disease conditions. Without this complexity, we risk missing important elements of these interactions. The endoplasmic reticulum (ER) stress response provides an opportunity to evaluate how GxE interactions alter gene expression across tissue type and stress conditions.

The ER is the largest cellular organelle and is a major site of protein and lipid synthesis, protein folding, and calcium storage. ER stress occurs when misfolded proteins accumulate in the ER lumen because of overwhelming protein folding demands or improper protein folding [2]. Cells respond to ER stress with the well conserved unfolded protein response (UPR), a coordinated series of cellular processes that reduces ER protein load and increases the ability of the ER to clear misfolded proteins [3, 4]. The UPR consists of changes in gene expression, reduction of protein translation, and degradation of misfolded proteins (ER associated degradation or ERAD). Gene expression changes are initiated through the three main signaling branches of the UPR: IRE1, ATF6, and PERK [5]. If the UPR is unable to restore ER homeostasis, it will activate apoptosis pathways. The response to misfolded proteins is essential for maintenance of basal cellular conditions and critical when there are any changes in the cellular environment, such as cellular differentiation and increased protein secretion [5].

Despite being a conserved, basic cellular function, the ER stress response is subject to inter-individual variation in *Drosophila*, mouse, and humans [6–9]. It is only recently that we are beginning to appreciate the impact of genetic variation on the ER stress response. In *Drosophila*, we used natural genetic variation to demonstrate that susceptibility to ER stress is highly variable across strains and is associated with SNPs in both canonical and novel ER stress genes [6, 7]. We also used mouse embryonic fibroblasts (MEFs) from inbred mouse strains to uncover a complex genetic architecture underlying the variable transcriptional response to ER stress among different genetic backgrounds [8]. Another study also demonstrated that the ER stress response is variable among immortalized human B cells from diverse individuals [9]. These studies nicely demonstrate a GxE effect, where different genotypes show vastly different transcriptional responses, dependent on the presence or absence of ER stress.

We previously showed that the highly variable ER stress transcriptional response in MEFs, derived from different genetic backgrounds, is driven by *cis*- and *trans*-regulatory variation [8]. There is an entire layer of genetic variation that remains silent under healthy conditions, but alters expression under ER stress. We also found that ER stress can have a profound effect on allele-specific expression (ASE) in both magnitude and direction. However, all past mouse and human studies examining the GxE interactions under ER stress utilized *in vitro* cell culture. Extensive GxE studies in other contexts have shown that genetic architecture changes drastically across different tissue types [1,10,11]. In this study, we addressed these limitations by mapping the genetic architecture underling GxE interactions during ER stress, *in vivo*, across tissues in the mouse.

Here we report that tissue type and ER stress have strong effects on how genetic variation impacts transcript levels. We identified hundreds of genes that showed variability in their response to ER stress that were dependent on tissue type and genetic background in mouse. These genes are enriched for processes, such as metabolism, inflammation, and gene expression. Strikingly, in contrast to previous studies where non-canonical ER stress genes were found to be variable,[8] we found that genotype-dependent ER stress response genes *in vivo* are enriched for terms with clear roles in the ER stress response, indicating that at least some variability in response is derived from canonical ER stress genes. Our study design employed F1 mouse crosses to uncover the *cis*- and *trans*-regulatory mechanisms that underlie the variable ER stress transcriptional response. We found a complex and context-specific regulatory landscape that underlies variability observed in the *in vivo* mouse. These regulatory mechanisms are context-specific in both number and strength. This study expands the understanding of the complex ER stress transcriptional response and the *cis*-/*trans*-regulatory mechanisms that impact this network in different environmental contexts. Together, these findings have implications for identifying ER stress response modifiers, better interpretation of regulatory mechanisms, and more comprehensive knowledge of the elements that make up the highly variable ER stress response.

## Results and Discussion

### *in vivo* ER stress induced by Tunicamycin

To evaluate the extent to which ER stress and tissue type affects gene expression in diverse genetic backgrounds, we induced ER stress in different strains of mice. We administered Tunicamycin (TM) or DMSO (control) with an intraperitoneal injection to mice from strains C57BL/6J (B6), CAST/EiJ (CAST), and their F1 progeny. TM induces ER stress by inhibiting protein N-glycosylation in the ER, causing an accumulation of misfolded proteins [12]. For this study, we focused on liver and kidney. An *XBP1* splicing assay and RT-qPCR of *BiP*, a canonical ER stress gene, show that TM injection induced a strong ER stress response (S1 Fig.). To determine the full transcriptional response to TM-induced ER stress, kidney and liver samples were analyzed by RNA-seq. ER stress-induced gene expression changes were identified by comparing control and TM samples in a tissue- and genotype-specific manner (S1 and S2 Tables). At a cutoff of 1.5-fold (5% FDR) change in transcript level, Gene Ontology (GO) enrichment analyses revealed enrichment for canonical ER stress response genes in all tissues and genotypes (S3 Table). Many canonical ER stress response genes, including *Hyou1, Hspa5* (*BiP*), *Ddit3* (*Chop*), and *Herpud1* [13–16] (Fig. 1), were significantly upregulated in both tissues and all three genotypes, indicating a strong UPR. As expected, injection of TM induces a robust *in vivo* ER stress-response.

**Fig 1.**
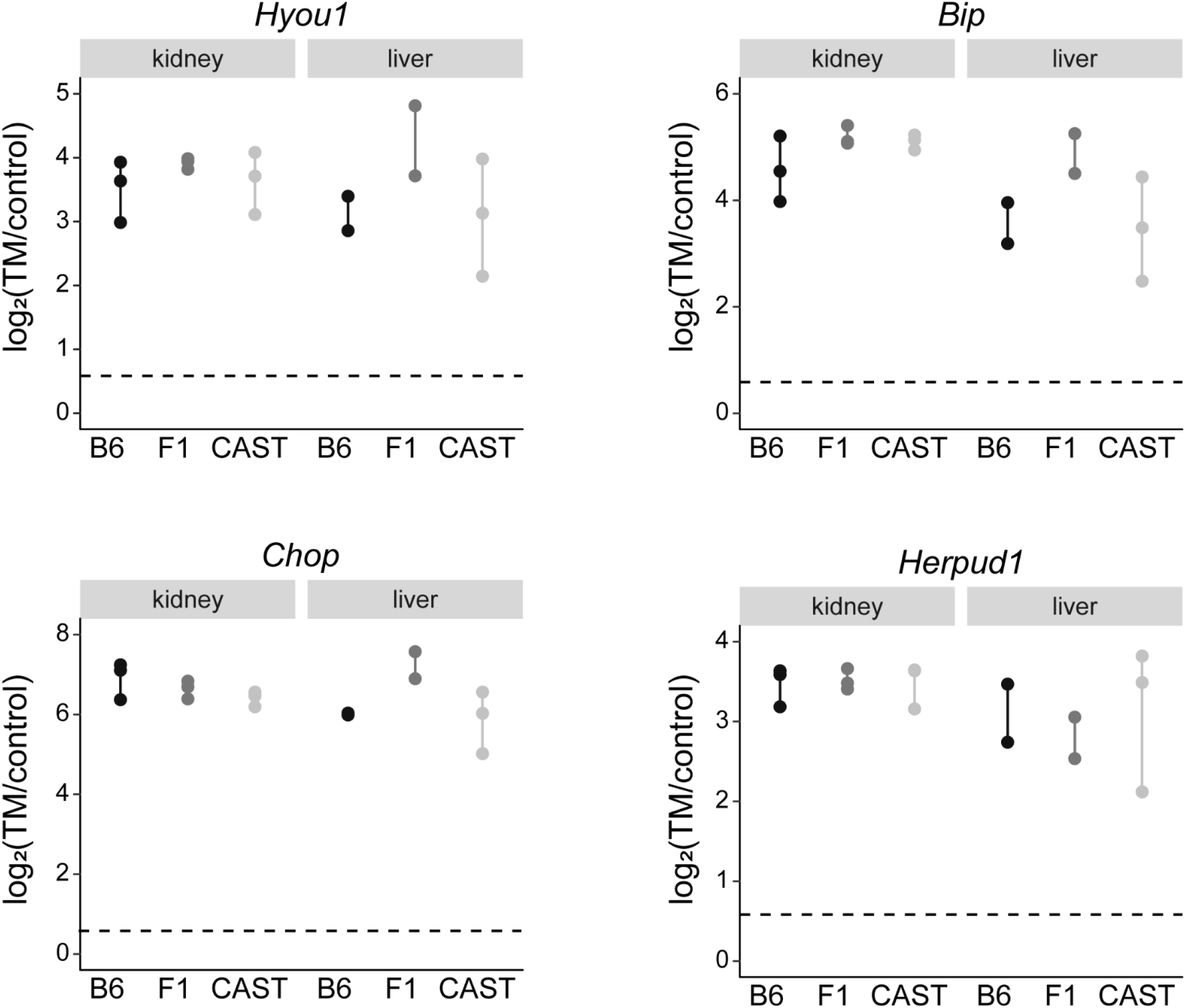
Upregulation of canonical ER stress genes across genotypes and tissues. Log_2_(TM/Control) is plotted for genes with known functions in the ER stress response. Dotted line indicates a 1.5-fold change in gene expression. Each point represents a biological replicate.

### ER stress-induced gene expression differs across tissues

We first characterized how the ER stress response differs by tissue type, irrespective of genotype. To do this we combined all three genotypes and compared only control and TM for liver and kidney samples. Of the 830 genes upregulated in liver, 264 (32%) were uniquely upregulated in liver (Fig. 2A). Of the 1,153 upregulated genes in kidney, 587 (49%) were unique to kidney. 566 genes were upregulated in both liver and kidney (68% in liver, 51% in kidney) (Fig. 2A). The upregulated genes shared between the liver and kidney were enriched for response to ER stress (GO:0034976, q= 2.80×10^-30^), ribosome biogenesis (GO:0042254, q=7.20×10^-23^), and ribonucleoprotein complex biogenesis (GO:0022613, q=1.70×10^-22^) (S4 Table). Of the 900 genes downregulated in liver, 794 (88%) were unique to liver. Of the 403 downregulated in kidney, 297 (74%) were unique to kidney. 106 downregulated genes were common to both liver and kidney (12% in liver, 26% in kidney) (Fig. 2B). The downregulated genes shared between the liver and kidney were enriched for oxidation-reduction process (GO:0055114, q= 3.90×10^-03^), single-organism metabolic process (GO:0008152, q= 3.90×10^-03^), and regulation of biological quality (GO:0065008, q= 1.30×10^-02^) (S4 Table). The proportion of common upregulated genes between the two tissues is much higher than common downregulated genes, suggesting that the upregulated response to ER stress is less dependent on tissue type (χ^2^, p<0.00001).

**Fig 2.**
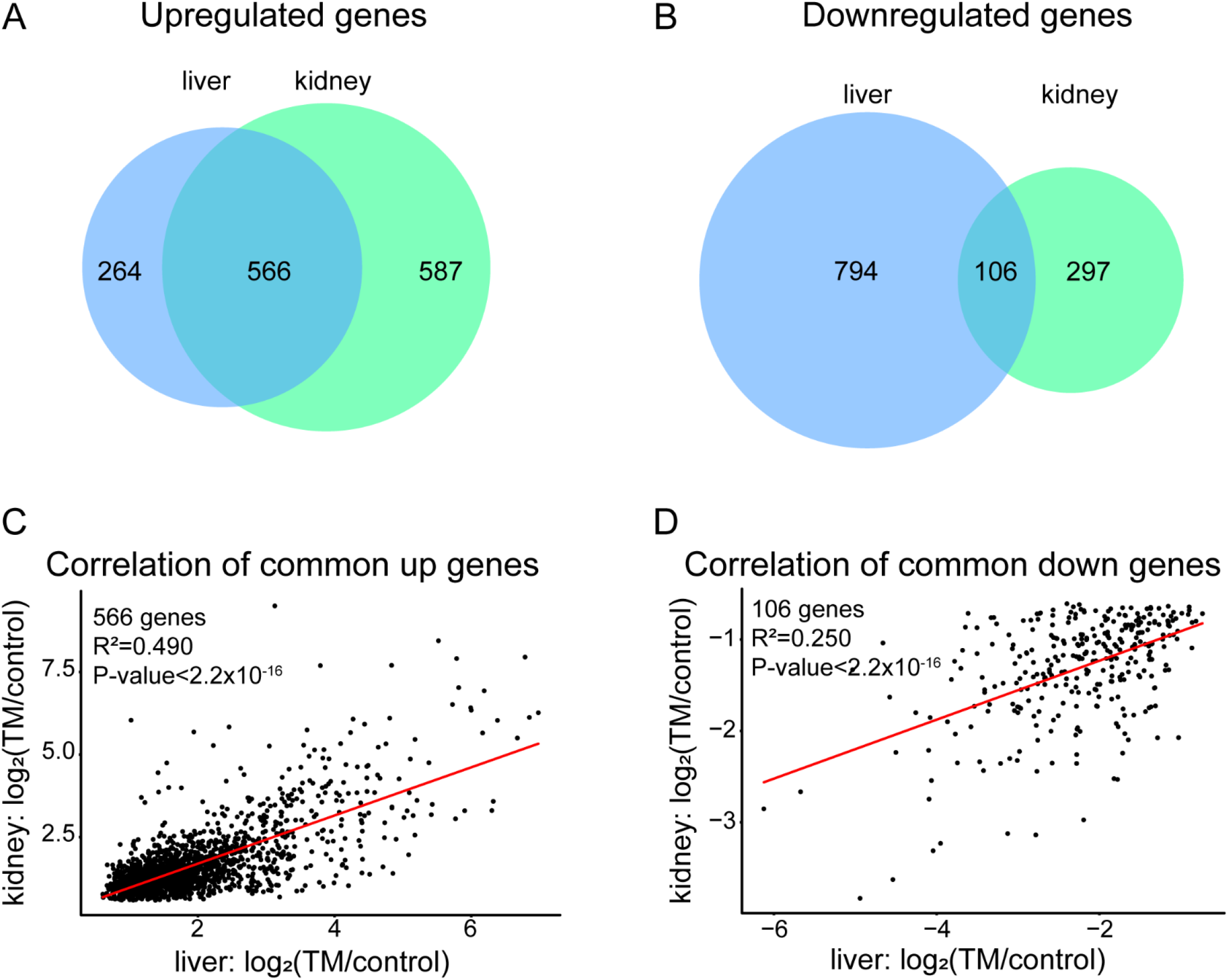
Tissue-specific ER stress-induced expression. Number of genes that display upregulation (A) or donwregulation (B) post ER stress in liver, kidney, or both tissues. C) Correlation of gene expression of the 566 common upregulated ER stress genes between liver and kidney. D) Correlation of gene expression of the 106 common downregulated ER stress genes between liver and kidney. Red line is regression line between the two tissues.

To examine whether common up- or downregulated genes in the two tissues differ in the magnitude of their response to ER stress, we performed a correlation analysis on the expression levels of common ER stress response genes. There is moderate correlation between liver and kidney expression of the 566 genes upregulated in both tissues (R^2^=0.490, p<2.2×10^-16^) (Fig. 2C). Despite the fact that these two tissues have very diverse functions, the common upregulated genes show similar expression levels in response to ER stress. There is a weaker correlation for the 106 genes commonly downregulated in both tissues (R^2^ =0.250, p<2.2×10-16) (Fig. 2D). A similar pattern emerges when correlations are examined for these overlapping response genes between tissues in the individual genotypes (S2 and S3 Figs.).

Next, we performed enrichment analysis on the 264 uniquely upregulated genes in liver and the 587 uniquely upregulated genes in kidney (S4 Table). The three most significantly enriched categories in liver were cellular metabolic process (GO:0044237, q= 4.9×10^-03^), organic cyclic compound metabolic process (GO:1901360, q=6.3×10^-03^), and nucleobase-containing compound metabolic process (GO:0006139, q= 6.6×10^-03^). In kidney, the three most significantly enriched categories were cellular macromolecule metabolic process (GO:0044260, q= 2.8×10^-15^), RNA metabolic process (GO:0016070, q= 7.4×10^-15^), and nucleic acid metabolic process (GO:0090304, q= 5.9×10^-12^). This enrichment indicates a metabolic response to ER stress that reflects the specific needs of each tissue. The 794 genes uniquely downregulated in liver were enriched for small molecule metabolic process (GO:0044281, q= 4.1×10^-68^), organic acid metabolic process (GO:0006082, q= 2.3×10^-64^), metabolic process (GO:0008152, q= 2.2×10^-63^), and other processes (S4 Table). We found no functional enrichment for the 297 genes that were uniquely downregulated in kidney.

While none of the most enriched tissue-specific processes are directly related to the canonical ER stress response, we did find individual canonical ER stress genes that respond in a tissue-specific manner. For example, *Creb3l2*, a known ER stress signal transducer,[17] is only significantly upregulated in liver and not kidney. Other genes uniquely upregulated in liver and known to be involved in the ER stress response or other stress responses include *Dnajc10, Erp29, Erp44*, and *Nrf1*. These genes are not responsive to ER stress in kidney. On the other hand, *Derl2*, which is involved in the degradation of misfolded proteins,[18] is uniquely upregulated in kidney. Other canonical ER stress genes uniquely upregulated in kidney include *Pmaip1, Cebpb, Nck1*, and *Niban1*. These genes are non-responsive to ER stress in liver. This demonstrates how even canonical ER stress genes can be differentially regulated in different tissues.

### ER stress-induced gene expression varies by genetic background

In order to identify how genetic background impacts variation in ER stress-induced gene expression, we utilized three genetically diverse genotypes (B6, CAST, and F1 hybrids). The majority of genes upregulated post-ER stress in either tissue are only upregulated in one or two of the genotypes based on the 1.5-fold cutoff (FDR 5%) (Fig. 3). In liver, 2,330 (of 20,131, 11.5%) genes are significantly upregulated post-ER stress in at least one of the genotypes. Of these genes, 950 (41%) are uniquely upregulated in only one genotype, 550 (23%) are upregulated in two genotypes, and 830 (36%) are upregulated in all three genotypes (Fig. 3). In kidney, 2,718 (of 20,661, 13.2%) genes are significantly upregulated post-ER stress in at least one of the genotypes. Of these genes, 971 (36%) are uniquely upregulated in only one genotype, 594 (22%) are upregulated in two genotypes, and 1153 (42%) are upregulated in all three genotypes (Fig. 3).

**Fig 3.**
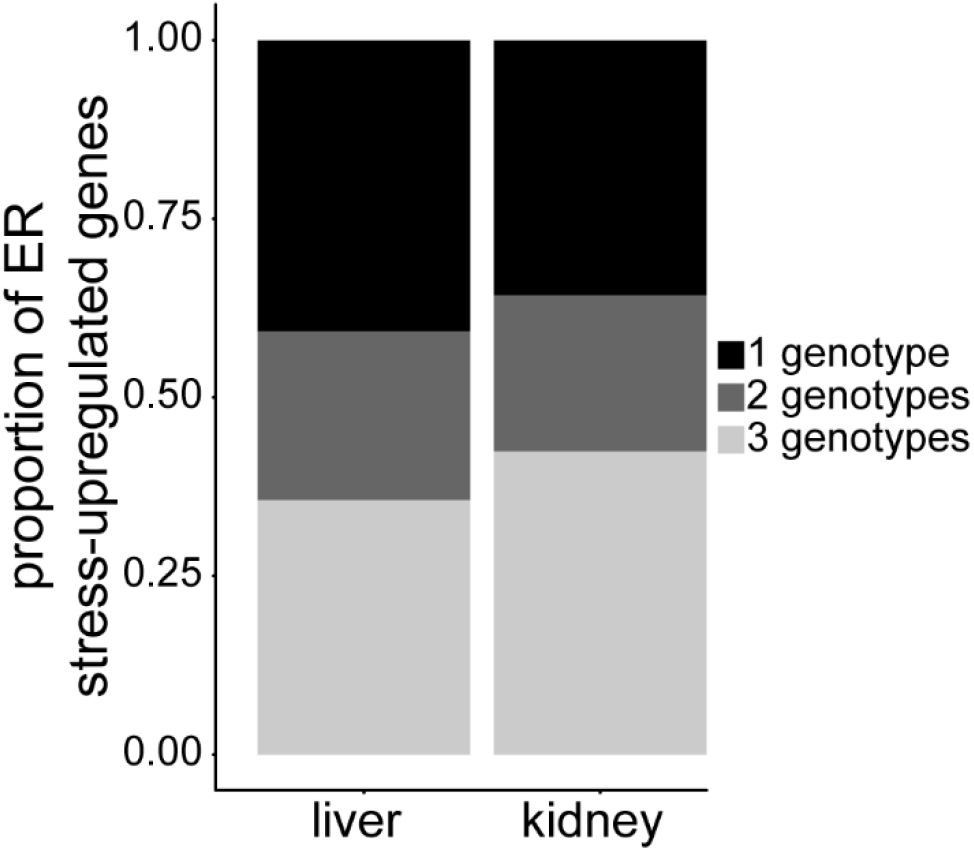
Genotype-specific expression post ER stress. Proportion of ER stress-upregulated genes that are shared in all three genotypes, shared in two genotypes, or unique to one genotype.

To identify genes in liver and kidney with the most variable, genotype-dependent ER stress-induced gene expression, we performed an ANOVA test to determine if there was an effect of genotype on expression for each tissue. For genes that are upregulated post-ER stress in at least one genotype in liver, 3.1% (72/2330; p ≤ 0.05) show variable expression levels, dependent on genotype (S5 Table). The three most significant GO categories for these genes were response to stress (GO:0006950, p=6.4×10^-4^), apoptotic signaling pathway (GO:0097190, p=1.0×10^-3^), and positive regulation of programmed cell death (GO:0043068, p=1.0×10^-3^) (S6 Table). In kidney, 21.4% (581/2718; p ≤ 0.05) of genes upregulated in at least one genotype are variable, dependent on genotype (S5 Table). The three most significant GO categories for these genes were ER to Golgi vesicle-mediated transport (GO:0006888, p=4.8×10^-10^), protein transport (GO:0015031, p=1.5×10^-9^), and protein folding (GO:0006457, p=1.3×10^-8^) (S6 Table). Only 8 genes display strain-dependent gene expression in both liver and kidney. The kidney had a significantly higher proportion of genes with genotype-dependent gene expression post-ER stress for upregulated genes (Liver: 3.1%; Kidney: 21.4%; χ^2^ p<0.0001). Tissue type has a large effect on how genetic background impacts gene expression.

The GO functional categories described above all encompass functions important to a proper ER stress response. A simple survey of these genes shows that many of them are canonical ER stress genes. For example, *Atf3* is a transcription factor that is activated by ER stress through the PERK pathway and contributes to the induction of other canonical ER stress factors [19]. *Atf3* expression is significantly variable in liver across the three genotypes (p=0.024). *Atf3* is significantly upregulated in all three genotypes, however, expression in CAST is significantly lower than the other two genotypes (Log_2_(TM/control): B6: 6.72, CAST: 4.54, F1: 7.66). In kidney, *Xbp1*, which is a major factor in the UPR, displays significant variable expression post-ER stress (p=0.012). *Xbp1* is significantly upregulated post-ER stress in all three genotypes; however, the degree to which *Xbp1* is upregulated in the F1 hybrid is significantly higher (Log_2_FoldChange: B6: 1.30, CAST: 1.36, F1: 2.17). Other highly variable genes include those not known to play a role in the ER stress response. One such gene is *Nupr1*, which displayed a significant genotype effect in both liver (p=0.011) and kidney (p=0.017) (Fig. 4). Nupr1 is involved in cell cycle regulation and autophagy. These processes are interconnected with the ER stress response [20, 21], providing a potential modifying role for *Nupr1* in the UPR. Other genes with no known role in the ER stress response that showed a genotype effects are highlighted in Figure 4. We observed genes where either the B6 or CAST parental genotype had increased or decreased expression, suggesting that the variation in gene expression upon ER stress is not solely driven by one strain.

**Fig 4.**
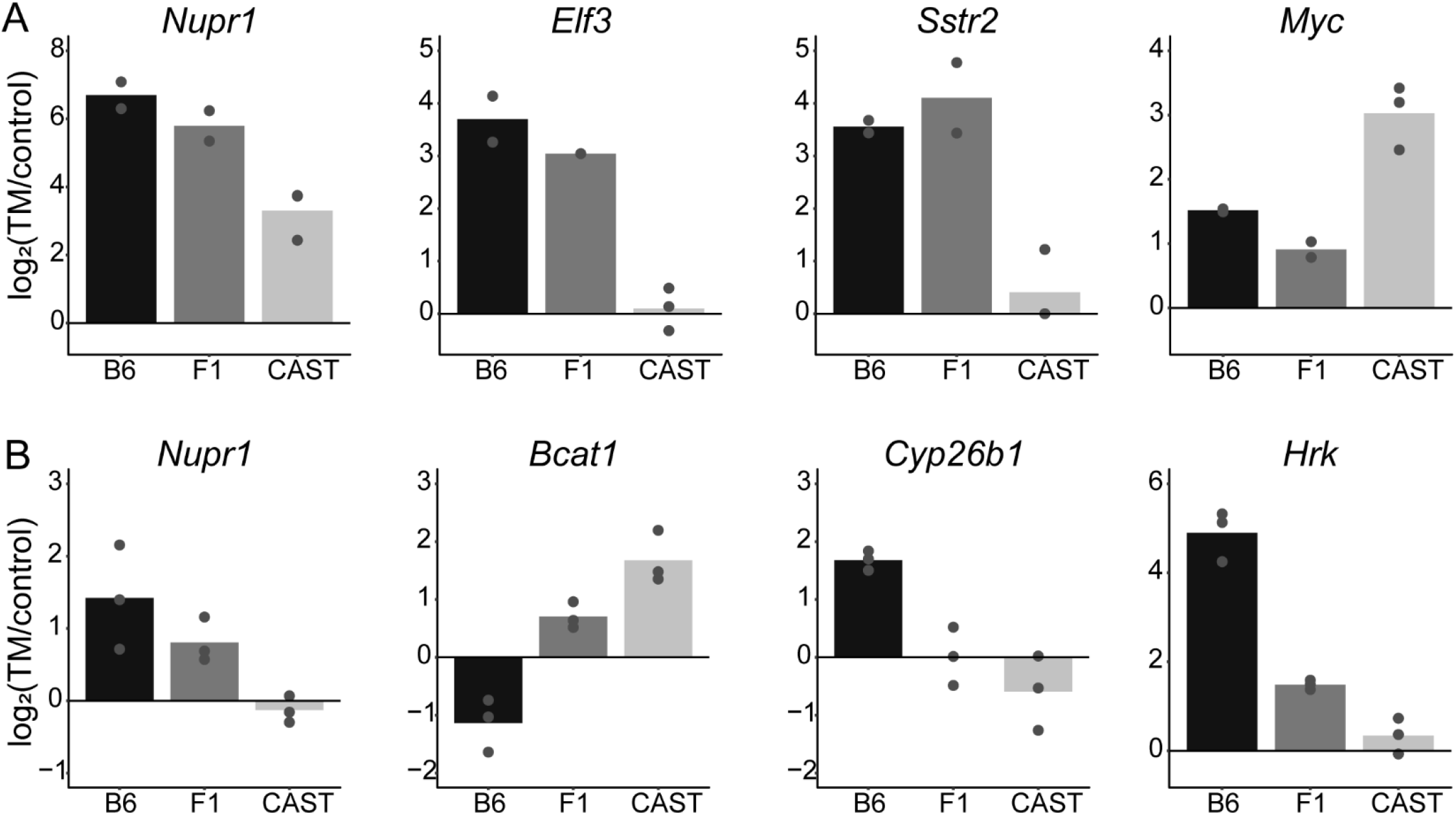
Variable expression of non-canonical ER stress genes across genotypes. Log2(TM/control) of counts for each genotype is plotted for a subset of non-canonical ER stress genes that displayed significant variable expression across the three different genotypes in liver (A) or kidney (B). Each point represents a biological replicate. All plots show a significant genotype effect (p<0.05).

Variable expression of transcription factors and other regulators of gene or protein expression can impact both canonical and non-canonical ER stress-responsive transcript levels [22]. In liver, a number of genes involved in transcription, such as *Atf3, Arntl, Elf3, Myc, Htatip2, Itga6*, and *Il23a*, show variable expression post-ER stress. As mentioned previously, *Atf3* shows variable expression and is involved in the UPR by activating canonical ER stress components. Another transcription factor and a non-canonical component of the UPR is *Arntl*, which is a critical component of the circadian clock. Dysregulation of circadian rhythm has been shown to have an impact on the UPR [23]. In kidney, there was an enrichment for genes involved in amino acid biosynthesis (GO:0008652, p=8.2×10^-8^) (S6 Table), such as *Bcat1, Bcat2, Cth, Asns, Pjgdh, Pycrl*, and *Psat1*. Regulation of amino acid synthesis is coupled with protein expression demands during ER stress [24]. The variable expression of amino acid biosynthesis genes could influence the availability of UPR upregulated proteins.

Next, we examined genotype effects on genes downregulated in response to ER stress. Again, we performed an ANOVA test on downregulated genes among the three genotypes. Of the genes that are downregulated post-ER stress in liver, 3.6% (101/2826, p≤0.05) show significant gene expression variability dependent on genetic background (S5 Table). The three most significant GO categories for these genes were cellular response to interferon-beta (GO:0035458, p=1.6×10^-5^), response to interferon-beta (GO:0035456, p=3.2×10^-5^), and defense response to other organism (GO:0098542, p=4.3×10^-4^) (S6 Table). The response to IFN-β stimulus has been shown to be tightly linked to the ER stress response [25]. One of these variably downregulated genes is *Ifit1*, which is involved in the innate immune system and inflammatory responses, responds to invasion by pathogens and endogenous damage signals [26]. *Ifit1* is downregulated approximately 4-fold stronger in the B6 liver (Log_2_(TM/control); B6: −2.576, CAST: − 0.523, F1: −0.728), suggesting that the liver immune and inflammatory response is subject to genetic variation post ER stress. For genes downregulated in kidney, 36.7% (772/2099; p≤0.05) show variability dependent on genotype (S5 Table). The three most significant GO categories for these genes were oxidation-reduction process (GO;0055114, p=2.5×10^-22^), metabolic process (GO:0008152, p=4.2×10^-18^), and lipid metabolic process (GO:0006629, p=1.9×10^-11^) (S6 Table). Given the connection between the ER stress response and these pathways [27, 28], any number of these oxidation-reduction genes could impact the ER stress response. Only 5 genes display downregulated strain-dependent gene expression in both liver and kidney. Again, compared to liver, kidney has a higher proportion of genes that display variable downregulation across the three genotypes (Liver: 3.6%; Kidney: 36.7%; χ^2^ p<0.0001). For both downregulated and upregulated genes, tissue type strongly influences the effect of genotype on gene expression post-ER stress. The incorporation of additional tissues will reveal the true extent to which genetic background can impact gene expression under stress conditions.

### Identification of *cis*- and *trans*-regulatory mechanisms

The identification of a highly variable ER stress response across different genetic backgrounds and tissues implicates a network of regulatory variation that impacts the expression levels of a wide range of genes, through either *cis*- or *trans*-mechanisms. A *cis*-regulatory variant influences the expression of a gene it is physically linked to. An example of a *cis*-mechanism is a promoter polymorphism impacting the gene’s expression levels. A *trans*-regulatory variant influences an unlinked gene, often physically distant from the variant. An example of a *trans*-mechanism is a polymorphism impacting a transcription factor that can then alter the expression of a wide range of genes across the genome.

To quantify allele-specific expression and partition the effects of genetic variation on gene expression into *cis*- and *trans*-mechanisms, we used F1 hybrid mice. Classification of *cis*- and *trans*-mechanisms was performed using previously published methodology [8, 29]. In order to determine if a gene is impacted by a *cis*- or *trans*-mechanism, we generated F1 hybrid mice by crossing the highly divergent parental strains B6 and CAST. Transcripts from F1 mice can be assigned to a parental chromosome based on parental SNPs in the transcript. *cis*- and *trans*-mechanisms for a particular transcript are assigned by comparing the ratio of allelic expression in the F1 to the ratio of total expression between the parental strains. In the F1 hybrid mouse, both parental alleles are exposed to the same *trans*-factors. Therefore, the ratio of allelic expression is a measure of *cis*-regulatory mechanisms between the two parental strains. If the allelic ratio matches the parental expression ratio, the expression difference is attributed to *cis*-mechanisms. If the allelic ratio differs from the parental expression ratio, the expression difference is attributed to *trans*-mechanisms.

We performed *cis-/trans*-analyses on liver and kidney under both control and TM conditions. Genes were assigned a *cis*- or *trans*-expression pattern or a combination of both mechanisms (FDR = 0.1%; S7-S9 Tables). For the remaining analyses, we focused on genes that displayed only a *cis*- or *trans*-mechanism. In liver, under control conditions, 580 transcripts displayed a *cis*-mechanism and 392 transcripts displayed a *trans*-mechanism (Fig. 5A). Under TM conditions, 617 transcripts displayed a *cis*-mechanisms and 449 transcripts displayed a *trans*-mechanism (Fig. 5B). In kidney, 710 transcripts displayed a *cis*-mechanism and 288 transcripts displayed a *trans*-mechanism under control conditions (Fig. 5C). Under TM conditions, 825 transcripts displayed a *cis*-mechanism and 230 transcripts displayed a *trans*-mechanism (Fig. 5D). The majority of the regulatory variation is due to *cis*-mechanisms.

**Fig 5.**
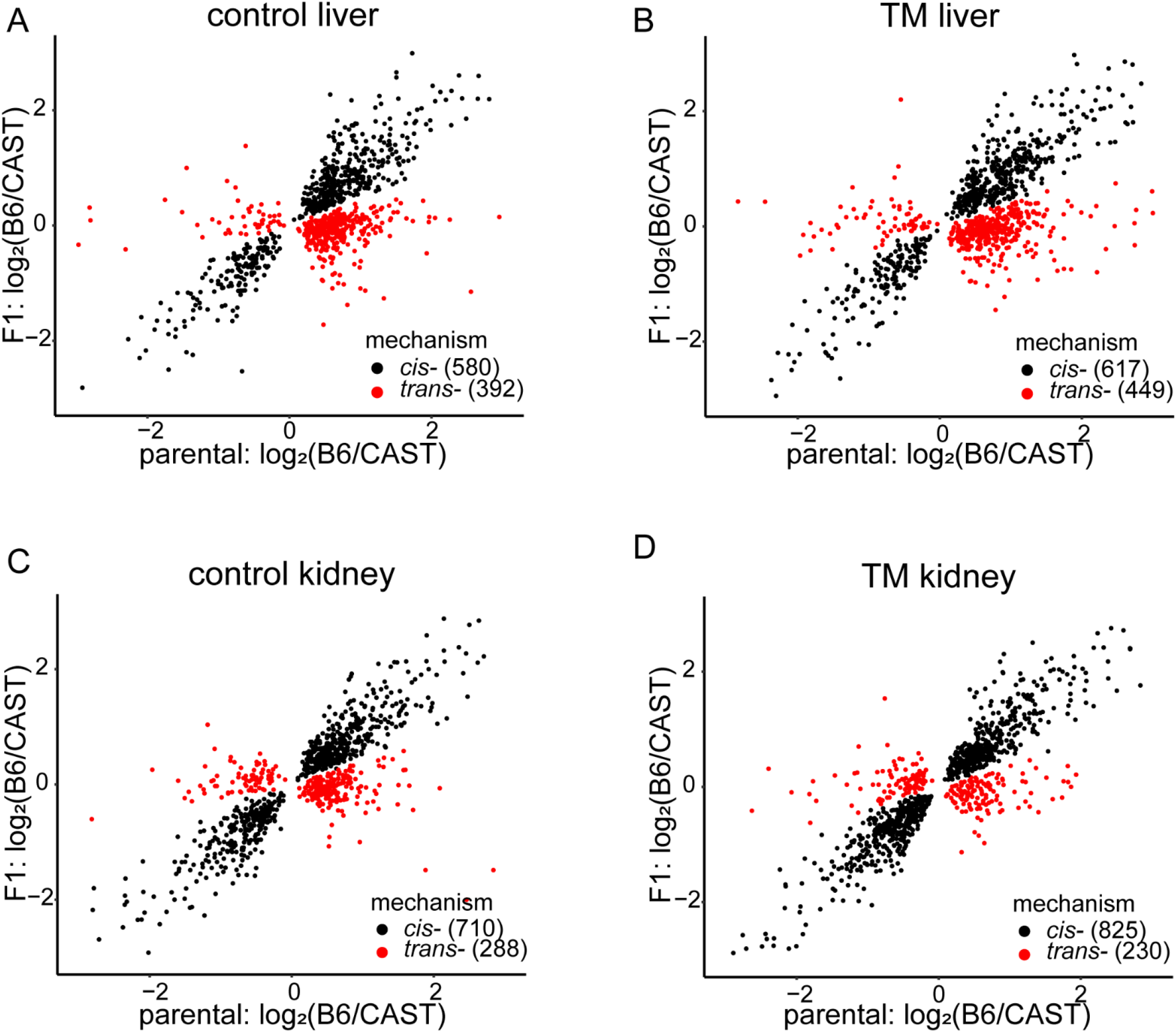
Expression ratios of genes with *cis*- or *trans*-mechanisms. Log2(B6/CAST) plotted for either alleles in the F1 hybrid or parental expression in control liver (A), TM liver (B), control kidney (C) and TM kidney (D). Each point represents a gene displaying either a *cis*- or *trans*-regulatory mechanism. Numbers in parenthesis are the number of genes that display that particular regulatory mechanism.

### ER stress reveals hidden regulatory variation unique to stress

To determine whether ER stress alters the contribution of *cis*- and *trans*-mechanisms to regulatory variation, we compared the proportion of transcripts displaying a *cis*- or *trans*-mechanisms in each tissue, under control and TM conditions. In liver, ER stress does not significantly alter the proportion of genes displaying a *cis*-mechanism (control: 0.59; TM: 0.58; P=0.413) (Fig. 6A). However, in kidney, there is a small, but significant increase in the proportion of genes with a *cis*-regulatory mechanism under ER stress conditions (control: 0.71; TM: 0.78; P=0.00024) (Fig. 6B).

**Fig 6.**
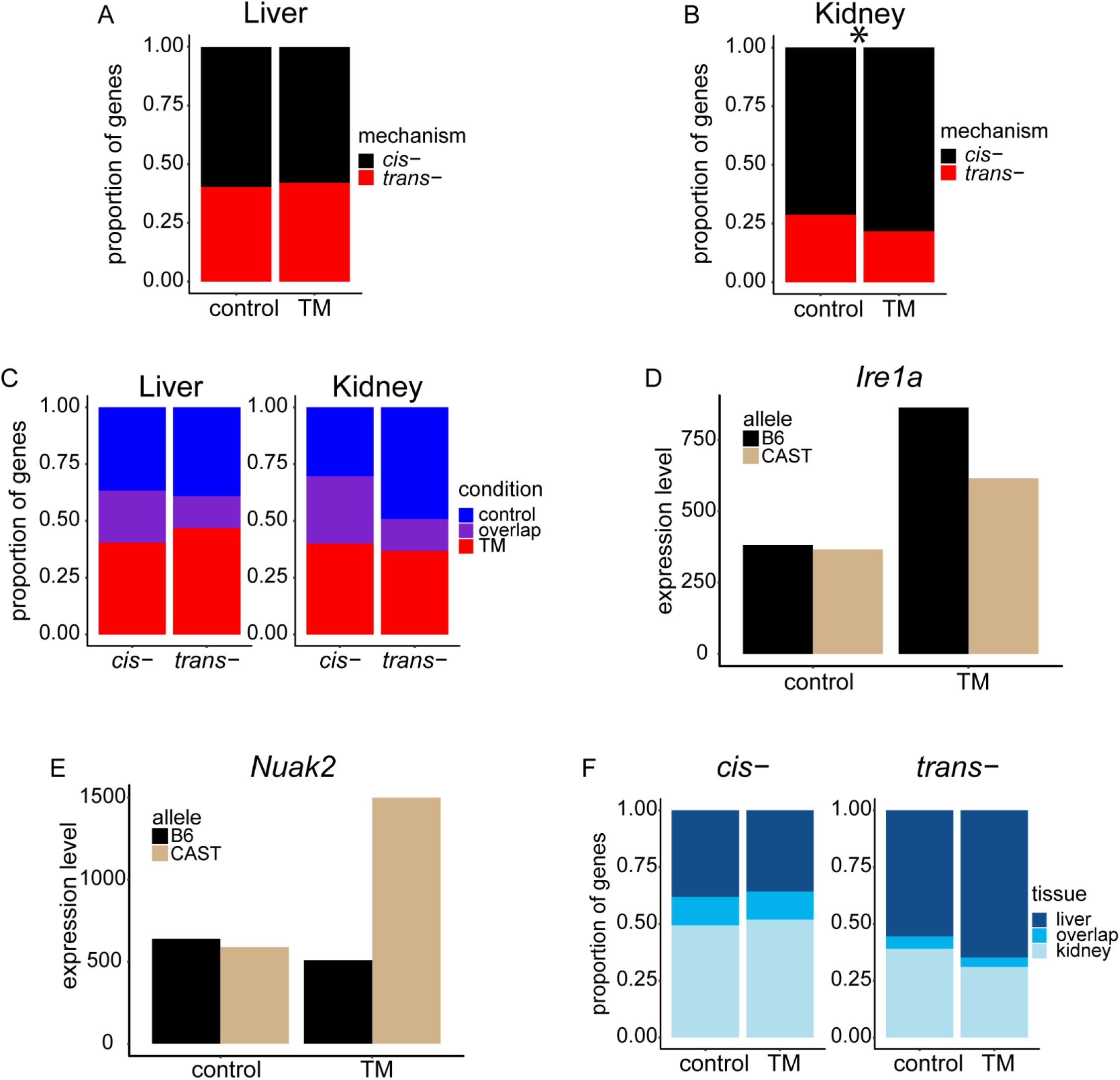
ER stress and tissue type reveal hidden regulatory variation. Proportion of genes displaying a *cis*- or *trans*-mechanism in liver (A) and kidney (B). Proportion of genes with either a *cis*- or *trans*-mechanism in only stress conditions, only control conditions or both, for liver and kidney (C). Examples of genes displaying *cis*-regulatory mechanisms only under stress conditions in F1 liver (D) and F1 kidney (E). Proportion of genes displaying a *cis*- or *trans*-mechanism seen in only liver, only kidney, or both (F). *P<0.00024

Because the ER stress transcriptional response involves hundreds of transcripts, it is likely that the actual genes showing *cis*- and *trans*-patterns are different under stress. To address this, the overlap under control and TM conditions, in both the liver and kidney were analyzed for genes showing *cis*- and *trans*-regulatory mechanisms. We observed both *cis*- and *trans*-regulatory mechanisms that were unique to control or stress conditions or present under both conditions.

In liver, of the *cis*-mechanisms detected under control or TM conditions, 37% (357/974) are unique to control, 40% (394/974) are unique to stress, and 23% (223/974) are common to both (Fig. 6C). Of the *trans*-mechanisms detected under control or TM conditions, 39% (288/737) are unique to control, 47% (345/737) are unique to stress, and 14% (104/737) are common to both. The magnitude of the common *cis*- or *trans*-mechanisms observed in both conditions were highly correlated (r =0.86, p<2.2×10-16) (S4A Fig.), suggesting that this common, overlapping regulatory variation is not impacted by ER stress.

In kidney, 30% (359/1183) of *cis*-mechanisms are unique to control, 40% (474/1183) are unique to stress, and 30% (351/1183) are common to both (Fig. 6C). Of the *trans*-mechanisms detected in kidney, 49% (224/454) are unique to control, 37% (166/454) are unique to stress, and 14% (64/454) are common to both. The magnitude of the common *cis*- or *trans*-mechanisms that were observed in both conditions were highly correlated (r^2^=0.91, p<2.2×10^-16^) (S4B Fig.) and likely not impacted by ER stress. The majority of genes that display a regulatory difference depend on the presence or absence of ER stress and those observed only in stress conditions may reveal critical components that might be responsible for the genotype-specific differences in the ER stress response.

In some cases, canonical UPR genes display regulatory variation only under stress conditions. For example, *Ire1α*, one of the main signal transducers of ER stress, displayed a strong *cis*-regulatory mechanism in the mouse liver that is only detectable under stress conditions (Fig. 6D). Under control conditions, *Ire1α* is expressed at similar levels by the B6 and CAST allele. Once ER stress is induced, the B6 allele is expressed 1.5-fold higher than the CAST allele. There are 691 SNPs within a ±2kb window of the *Ire1α* gene that differs between the B6 and CAST genotype. Any one or a combination of these SNPs could be contributing to this *cis*-regulatory difference. Genes that have not been implicated in the ER stress response, but show a strong, stress-specific regulatory difference, might represent novel UPR genes and pathways. For example, in the mouse kidney, the gene *Nuak2* displayed a strong *cis*-mechanism seen only under stress conditions (Fig. 6E). While under control conditions the *Nuak2* alleles are equally expressed, but in the stressed F1 mouse kidney, the CAST allele is expressed 3-fold higher than the B6 allele. *Nuak2* belongs to the AMPK protein kinase family and has mainly been linked to cancer [30, 31]. Understanding *Nuak2* and its role in AMPK regulation and ER stress can provide insight into the role of *Nuak2* in human disease such as cancer.

### ER stress reveals hidden regulatory variation unique to tissue type

We next asked whether there were tissue-specific regulatory mechanisms. We investigated the overlap of genes that displayed *cis*- or *trans*-regulatory mechanisms between liver and kidney in control and TM conditions. Of the genes that display a *cis*-regulatory mechanism under control conditions, 38% (436/1,146) are unique to liver, 49% (566/1,146) are unique to kidney, and 13% (144/1,146) are common to both (Fig. 6F). Of the genes that display a *cis*-regulatory mechanism under TM conditions, 36% (458/1,283) are unique to liver, 52% (666/1,283) are unique to kidney, and 12% (159/1,283) are common to both (Fig. 6F). The magnitude of the *cis*-mechanisms observed in both tissues were moderately correlated (Control: r^2^=0.425, p<2.2×10^-16^; TM: r^2^=0.408, p<2.2×10^-16^) (S5A and S5B Figs.).

Of the genes that displayed a *trans*-regulatory mechanism under control conditions, 55% (357/644) are unique to liver, 39% (252/644) are unique to kidney, and 6% (35/644) are common to both (Fig. 6F). Of the genes that display a *trans*-regulatory mechanism under TM conditions, 65% (422/652) are unique to liver, 31% (203/652) are unique to kidney, and 4% (27/652) are common to both (Fig. 6F). The magnitude of the *trans*-mechanisms observed in both tissues showed a small correlation only in control conditions (Control: r^2^=0.186, p=0.009; TM: r^2^=0.126, p=0.069) (S5C and S5D Figs.). Under control and TM conditions, we found that more genes with a *cis*-mechanism were common between the two tissues than genes with a *trans*-mechanism (Control: χ^2^ p < 0.00001; TM: χ^2^ p < 0.00001).

Under TM conditions, the majority of genes that displayed a regulatory mechanism were unique to either liver or kidney. Any one of these genes with a tissue- and stress-specific regulatory mechanism could be a gene involved in inter-individual variation in tissue-specific ER stress responses. For example, the genes *Lama5, Hnf4a, Scnn1b*, and *Pkd2* all display strong *cis*-regulatory mechanisms under stress conditions in kidney that are not observed in the mouse liver. Each of these genes were previously implicated in kidney diseases, such as Liddle’s syndrome and Polycystic kidney disease [32–35]. The kidney is a tissue that relies heavily on protein transport and secretion. Proper ER function and response to ER stress plays a large part in kidney function. In fact, ER stress and aberrant protein trafficking is pathogenic in a large number of kidney diseases [36]. In liver, for genes such as *Sidt2* and *Adk*, they display *cis*-regulatory mechanisms that are unique to the stressed liver and have been associated with human fatty liver disease [37, 38]. These genes with tissue-specific *cis*-regulatory mechanisms are clear examples of how genetic variation has a differential impact across tissues.

### Magnitude of effect of regulatory mechanisms on gene expression

To determine the magnitude of the effect of *cis*- and *trans*-regulatory mechanisms on gene expression levels, we compared the ratio of the absolute fold change of the parental B6 expression to the parental CAST expression for each gene displaying a regulatory mechanism. In liver, under control and TM conditions, *cis*-regulatory differences have a stronger effect on gene expression levels than *trans*-regulatory differences (control: P<0.003; TM: P<1.6×10^-5^) (Fig. 7A). There was no difference in the magnitude of effect when comparing *cis*- or *trans*-regulatory mechanisms across control and TM conditions in liver (*cis*-: P=0.26; *trans*-: P=0.93) (Fig. 7A). We found a similar pattern in kidney. *cis*-regulatory differences have a stronger effect on gene expression levels than *trans*-regulatory differences in control and TM conditions (control: P<9.0×10^-7^; TM: P<7.06×10^-4^) (Fig. 7B). Again, there was no difference in the effect when comparing *cis*- or *trans*-regulatory mechanisms across conditions in kidney (*cis*-P=0.25; *trans*-P=0.98) (Fig. 7B).

**Fig 7.**
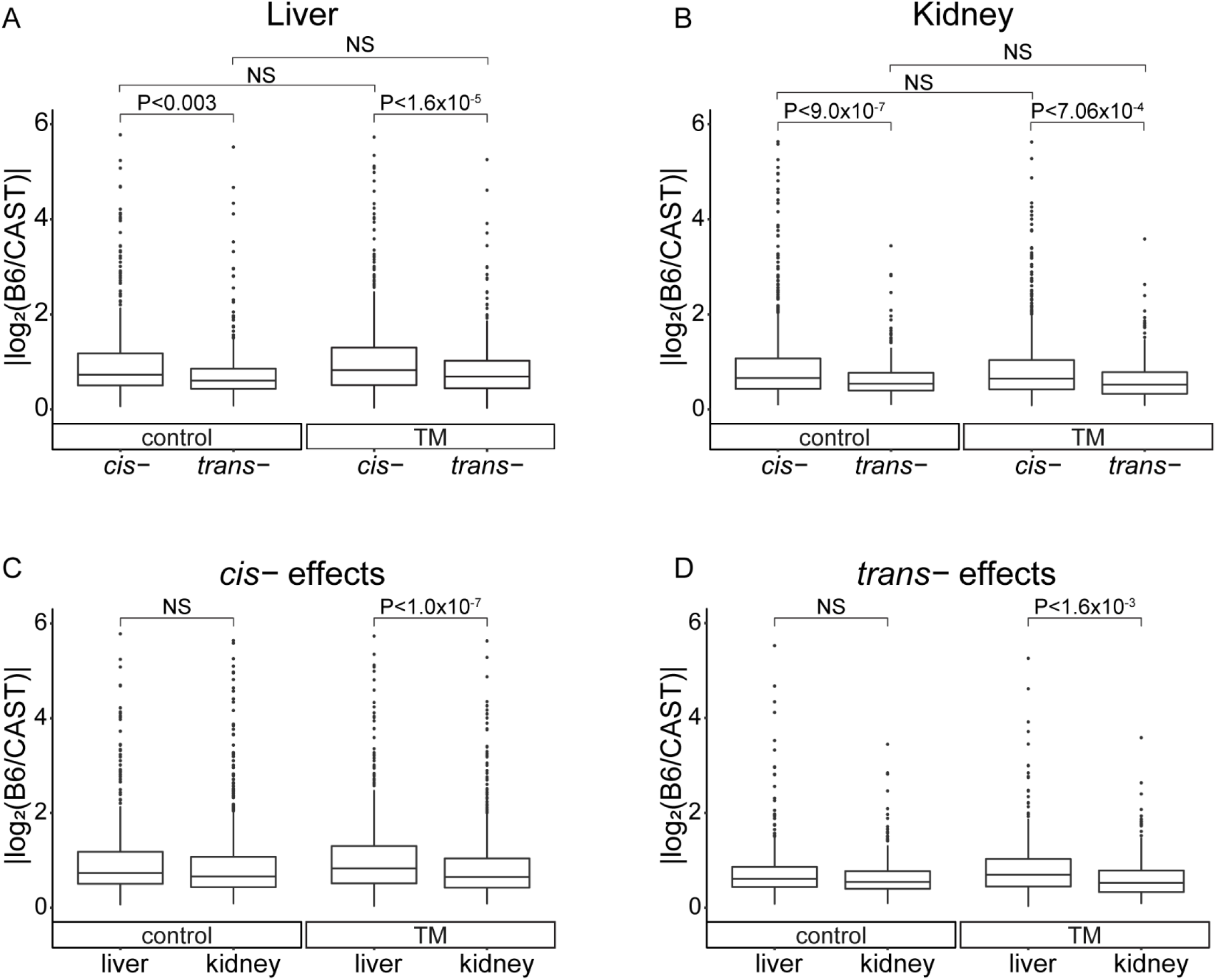
Impact of *cis*- and *trans*-mechanisms on gene expression. Absolute log2 fold change of parental expression for genes displaying a regulatory mechanism. Comparing genes with a *cis*- or *trans*-mechanism in either control or stress conditions in liver (A) or kidney (B). Comparing impact of *cis*-mechanisms (C) or *trans*-mechanisms (D) on gene expression in control or stress conditions. Liver: control *cis*-: mean=1.04, SD=1.03 | control *trans*-: mean=0.82, SD=0.91 | TM *cis*-: mean=1.14, SD=1.09 | TM *trans*-: mean=0.86, SD=0.74; Kidney: control *cis*-: mean=0.94, SD=0.89 | control *trans*-: mean=0.66, SD=0.44 | TM *cis*-: mean=0.87, SD=0.74 | TM *trans*-:mean=0.65, SD=0.48. NS= not significant.

We compared the effects of *cis*- and *trans*-mechanisms between liver and kidney to better understand the impact of tissue type on the strength of a regulatory mechanism. Under control conditions, there is no difference between the strength of *cis*-mechanisms between the two tissues (P=0.170). However, under TM conditions, *cis*-mechanisms in liver have a stronger effect on gene expression than in kidney (P<1.0×10^-7^) (Fig. 7C). *trans*-mechanisms, under control conditions, also showed no difference between tissues, but were stronger in liver than kidney under TM conditions (control: P=0.20; TM: P<1.6×10^-3^) (Fig. 7D). Within a condition, *cis*-mechanisms on average are stronger than *trans*-mechanisms, but tissue type and ER stress have different effects on the strength of regulatory mechanisms. Genetic variation within the stressed liver has a stronger effect on transcription than in any other environmental combinations.

### ER stress induced change in allele-specific expression

Next, we tested whether ER stress and tissue type altered allele-specific expression (ASE). ASE is measured as the ratio of allelic expression in the F1 mouse (B6/CAST allele). We compared the allelic ratio in the F1 under stress and control conditions, using a Fisher’s exact test (5% FDR). A significant change in ASE post-ER stress was observed in 17% and 13% of expressed transcripts in liver (970/5669) and kidney (1010/7764), respectively (Fig. 8A and C) (S10 Table). The liver displayed a higher proportion of genes with a change in ASE under stress (χ^2^; p=0.00001). Change in ASE in both tissues was driven equally by CAST and B6 alleles, indicating that there is no unexpected hybrid effect (S6A Fig.).

**Fig 8.**
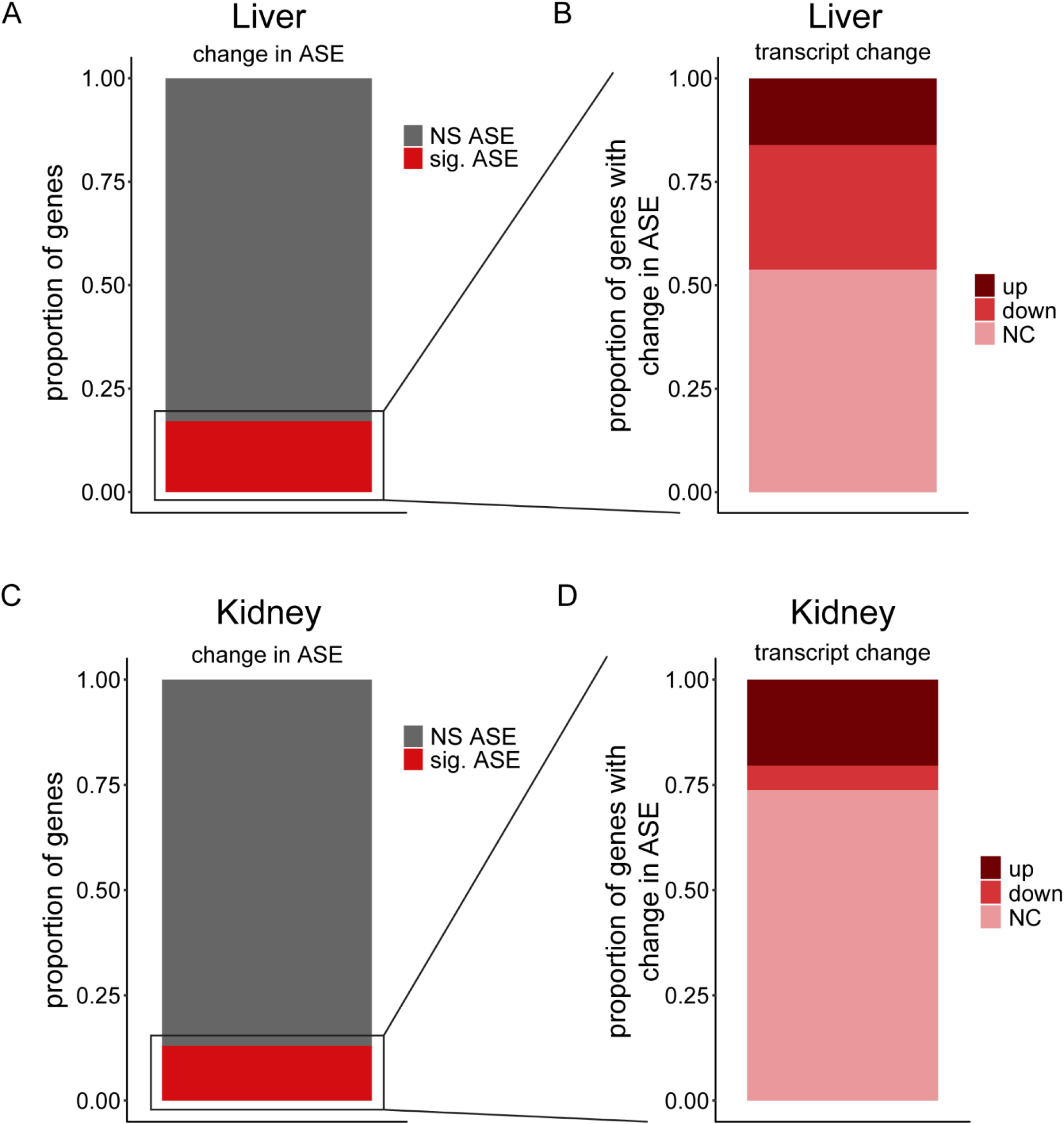
ASE and corresponding change in RNA transcript. Proportion of genes in the F1 displaying significant change in ASE in liver (A) and kidney (C). Proportion of genes with a significant change in ASE showing ER stress-induced increase in RNA transcripts, decrease, or no change in liver (B) and kidney (D).

We next asked if genes with changes in ASE post-ER stress also show ER stress-responsive transcript levels. The majority of genes with a change in ASE post-ER stress do not have ER stress-responsive transcripts (liver: 522/970 or 54%; kidney: 745/1010 or 74%) (Fig. 8B and D). Genes with a change in ASE and transcript level post ER stress fall in both up- and downregulated categories (Liver: 35% upregulated 156/448, 65% downregulated 292/448; Kidney: 78% upregulated 206/265, 22% downregulated 59/265) (Fig. 8B and D) In all cases, the B6 and CAST alleles contributed equally to changes in ASE (S6C and S6D Figs.).

Genes that show a significant change in ASE and in transcript levels post-ER stress are of particular interest, as this suggests differential allelic response to stress. *Sesn2*, which is involved in a variety of different stress responses,[39] showed one of the most significant changes in ASE (Fisher’s exact; q<0.00001). Under control conditions, the B6 and CAST allele in the F1 hybrid mouse are expressed at equal levels (B6: 0.53; CAST: 0.47). Total *Sesn2* transcript responded to TM conditions with a 13-fold increase. At the allelic level, the CAST allele is increased 24-fold, while the B6 allele is only increased 3-fold (TM: B6: 0.13, CAST: 0.87) (Fig. 9A). The large increase of the CAST allele is driving the *Sesn2* transcriptional response to TM-induced ER stress in the F1. This strong allelic response indicates that *Sesn2* contains a strain- and ER stress-specific *cis*-element that drives differential transcriptional response to ER stress and potentially other stress stimuli such as hypoxia and reactive oxygen species [39]. We see similar patterns for genes that are downregulated post-ER stress. *Presenilin 2* (*Psen2*),which cleaves proteins such as APP (amyloid-beta precursor protein) [40] and has been shown to cause Alzheimer’s Disease, displays a change in ASE (Fisher’s exact; q=0.00001) and a 2.7-fold decrease in RNA transcript levels (Fig. 9B). Under control conditions, the B6 allele is more highly expressed (B6: 0.72; CAST: 0.28). Under stress conditions, the *Psen2* B6 allele decreases 4.2-fold in expression while the CAST allele only decreases 1.6-fold. The drastic reduction of the B6 allele results in near equal expression levels of the two alleles under stress conditions (B6: 0.47; CAST: 0.53).

**Fig 9.**
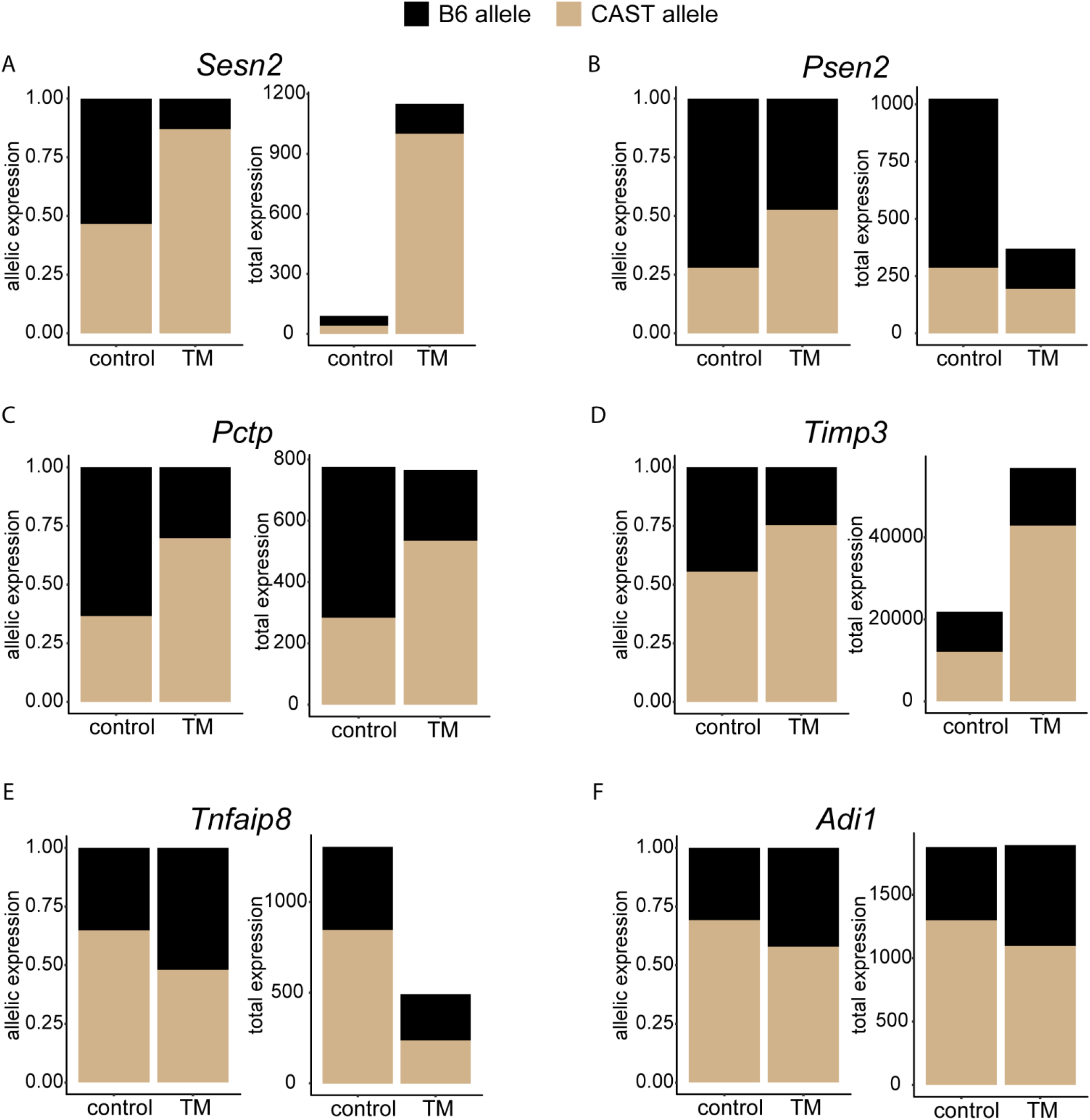
Change in ASE and transcript levels across stress conditions. ASE and total RNA expression levels are plotted. Genes that display ER stress-induced change in ASE in liver (A-C) or kidney (D-F). (A) and (D) show an increase in total transcript levels, (B) and (E) show a decrease in total transcript level, and (C) and (F) show no change in transcript levels.

As mentioned above, the majority of genes that have a change in ASE post-ER stress do not have significant changes in transcript levels. Phosphatidylcholine transfer protein (*Pctp*) shows a significant change in ASE, but no change in RNA transcript levels (Fig. 9C). Under control conditions, the B6 allele accounts for 63% of allelic expression, while under TM conditions, the B6 allele accounts for only 30% of allelic expression (Fisher’s exact; q=0.00001). While the ratio of expression between alleles is significantly changed under TM conditions, the net result is no significant change in total RNA transcript levels, suggesting a possible compensatory mechanism between the two alleles. Similar patterns emerge in kidney with a wide range of genes (Fig. 9D-F).

The majority of ASE post ER stress is tissue-specific. Only 210 transcripts display a change in ASE post-ER stress common to both tissues (Liver: 210/970, 22%; Kidney: 210/1010, 21%) (S7 Fig.). For these common genes, tissue type had a strong effect on the magnitude of the ASE changes post ER stress, in line with what we observed with tissue-specific changes in magnitude of *cis*- and *trans*-mechanisms. For example, in *Cathepsin L* (*Ctsl*) (Fig. 10A), the CAST allele is more ER stress responsive in kidney, while in liver, the B6 allele is more responsive. *Ctsl*, which is involved in lysosomal protein degradation, is upregulated in both liver (FC= 4.3) and kidney (FC= 3.2) post-ER stress. *Ctsl* also shows a change in ASE in liver (q=0.017) and kidney (q=0.0002). However, in liver, the CAST allele was responsible for only 55%of the increase in expression levels, but in kidney, the CAST allele was responsible for 71% of the increase in expression levels. This pattern was also observed in downregulated genes. *Flavin containing dimethylaniline monoxygenase 1* (*Fmo1*) (Fig. 10B), which is involved in the oxidation reduction process, is downregulated in both liver (FC=-3.54) and kidney (FC=-1.5) and shows a change in ASE in liver (q=7×10^-5^) and kidney (q<0.00001). The CAST allele in liver accounts for 80% of the expression downregulation, but only 28% of the downregulation in kidney.

**Fig 10.**
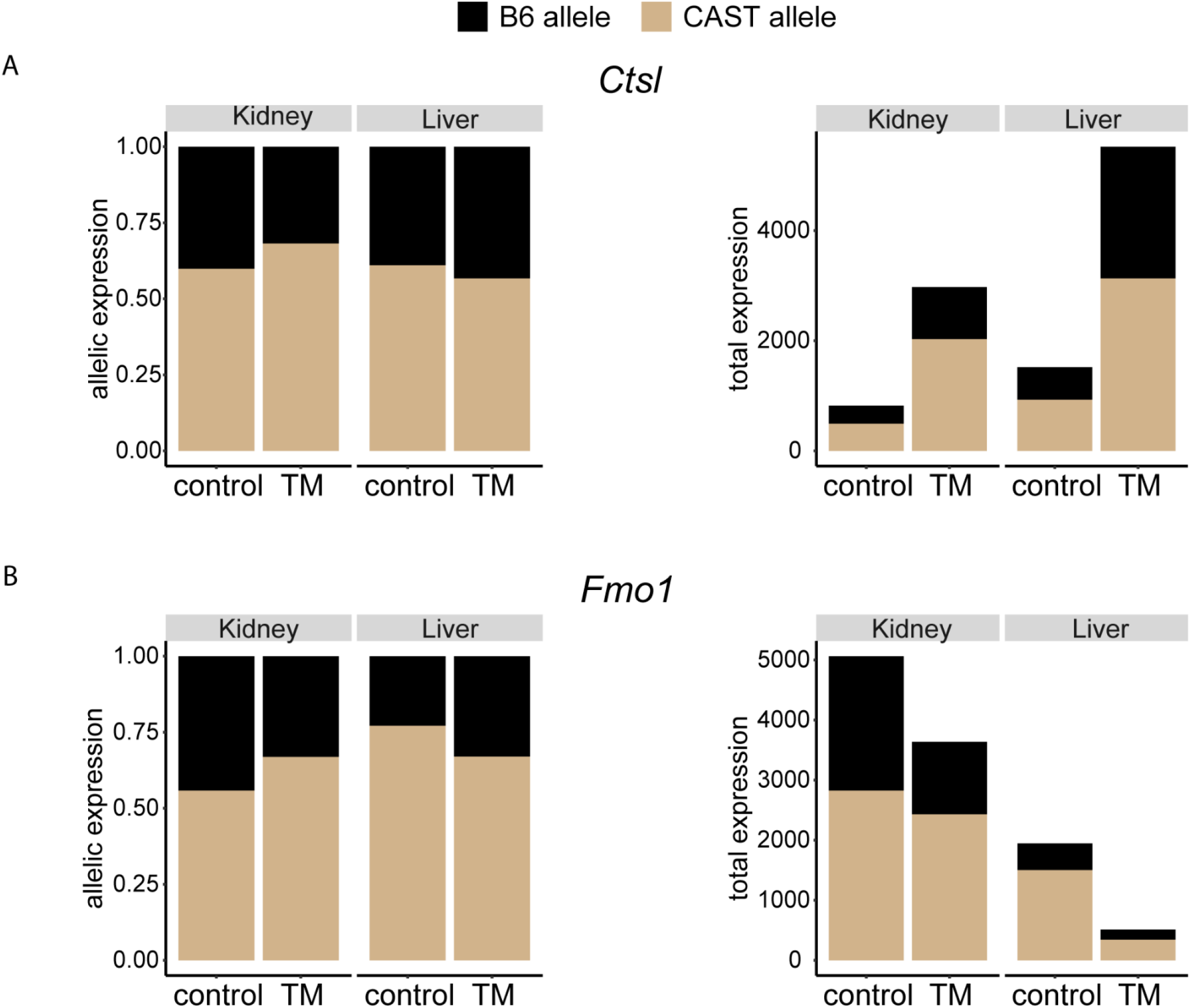
Variable magnitude in ASE across tissues. Examples of genes that show ASE in both tissues, but differ in the magnitude of that ASE in a tissue-dependent manner. An example of an upregulated gene (A) and a downregulated gene (B). ASE and total RNA expression levels are plotted.

### Transcription factor binding site enrichment in common and variable ER stress genes

The large transcriptional response to ER stress is partly driven by three major transcription factors, XBP1, ATF6, and ATF4, binding in *cis*-near a gene. While we cannot identify them with this study design, we hypothesized that at least some of the *cis*-regulatory variation sits within these transcription factor binding sites. We searched for enrichment of these sites near ER stress responsive genes from this study (S1 and S5 Tables). First, we identified TF binding site enrichment in the ER stress responsive genes common to all genotypes and tissues (S1 and S11 Tables). Some transcription factor binding sites are associated with canonical ER stress response genes. Two of these sites, both bound in conjunction by NFYA and ATF6, are called the ER Stress Response Elements I and II (ERSEI and ERSEII) [41, 42]. Within these common genes, NFYA binding sites were most significantly enriched (Z-score = 54.8). There was also enrichment for CEBPA binding sites (Z-score = 3.4), which can bind in conjunction with CHOP [43], a downstream component of the UPR involved in ER stress induced apoptosis signaling [44]. CHOP has also been implicated in regulation of metabolism, especially in the mouse liver, and inflammation [43–46]. We also observed enrichment for KLF4 binding sites, a transcription factor involved in proliferation and apoptosis (Z-score = 24.3) [47]. HIF1A binding sites were also highly enriched (Z-score = 18.5). HIF1A is a hypoxia induced transcription factor that is involved in oxygen homeostasis [48]. There is significant overlap in genes between the hypoxic response pathways and the ER stress response [49]. We also see significant enrichment for Nf-kappaB which is involved in immunity and inflammation (Z-score = 3.2) [50].

Next, we analyzed the set of variable ER stress responsive genes that showed genotype-dependent expression post-ER stress (S5 and S11 Tables). For genes variable in liver, we found enrichment for the CTCF binding site (Z-score = 11.6) near these variable ER stress response genes. CTCF is involved in the regulation of chromatin which can influence an even wider range of genes and their expression levels. Other significant binding sites for genes variable in kidney include ARNT (Z-score = 8.9) and PAX4 (Z-score = 9.8), which are involved in leukemia and development respectively. Similar to the common ER stress responsive genes, there is also enrichment for genes involved in immunity and inflammation, like RELA and Nf-kappaB (Liver: Z-score = 6.4 and 5.6; Kidney: Z-score = 3.6 and 4.5), mirroring the results found in MEFs [8].

### Conclusions

The ER stress and large UPR transcriptional response provides a unique opportunity to study how GxE interactions can alter gene expression levels. We took advantage of two genetically diverse mouse strains, B6 and CAST, and their F1, and induced a strong *in vivo* ER stress transcriptional response. This provided the opportunity to study how stress and tissue type impacts the effect of genetic variation. We uncovered genes that showed variable transcript levels in a genotype x tissue x stress manner. These genes implicated networks and pathways that could contribute to the variable ER stress response. Additionally, the F1 hybrid gave us the ability to uncover the regulatory mechanisms that are impacted by stress and tissue type. We discovered that most *cis*- and *trans*-regulatory mechanisms are context specific, with most unique to only one context. Altogether our results provide a better understating for how genetic background and tissue type impacts the genetic architecture of the inter-individual transcriptional response to ER stress in mouse and how different genotypes respond to different environments

We previously used mouse embryonic fibroblasts (MEFs) to assay how a complex genetic architecture influences the transcriptional response to ER stress across different genetic backgrounds [8]. Here, we utilized an *in vivo* mouse model to identify how these patterns change when comparing across different tissues. In the MEF study, upregulated genes most significantly influenced by genetic background did not have canonical ER stress functions [8]. However, in this current study, we found enrichment for functions such as response to stress, protein transport, and protein folding, all canonical ER stress functions. In contrast to the MEF study, this work demonstrates that the genotype-dependent transcripts upregulated in liver and kidney have clear roles in the UPR. Much of the variability in the response to ER stress likely stems from these genes. Interestingly, in this current study, we found enrichment for inflammation in the context of genotype–dependent downregulated genes, particularly in liver. This is in opposition to the MEF study which found enrichment for inflammation in upregulated genes [8]. This highlights the different ways that genotype interacts with the inflammatory response during the ER stress response, especially in different tissues.

This study emphasizes the strong effect that tissue type has on how genetic variation impacts the transcriptional response to ER stress. These differences are observed in the number of regulatory mechanisms and the strength of these mechanisms across tissues. The ER stress response is implicated in many different diseases such as Alzheimer’s disease, type II diabetes, ALS, atherosclerosis, and cancer [51–56]. Each of these diseases are unique in their tissue of origin. Additionally, each individual diagnosed with an ER stress-related disease will have different genetic backgrounds. The tissue-dependent ER stress response observed in this study illustrates how future studies involving ER stress and disease should investigate the disease-relevant tissue in the context of different genetic backgrounds, potentially uncovering novel tissue-specific effects and mechanisms.

This study utilized genetic variation present in different strains of mice to demonstrate the strong impact that genetic background and tissue type has on cellular processes such as the ER stress response. We detected numerous ER stress- and tissue-specific responses in expression levels, regulatory mechanisms, regulatory mechanism strengths, and allele-specific effects. The majority of these findings would have been missed if only studying one genotype, condition, or tissue. Future studies can reveal even more complex interactions that affect transcriptional levels by incorporating more environmental variables, such as cell type, additional tissues, and other cellular stressors. This type of analysis provides better predictive power for how a variant will impact expression across different environments.

## Materials and Methods

### Mice

C57BL/6J (B6) and CAST/EiJ (CAST) mice were obtained from Jackson Laboratories (Bar Harbor, ME). F1s were generated by crossing female B6 mice to male CAST mice. Parental strains were generated by crossing B6 mice to B6 mice and CAST mice to CAST mice. 6 male mice of each genotype (B6, CAST, and F1) of approximately 15-23 weeks of age were used for the experiment. All experiments involving mice were performed according to institutional IACUC and NIH guidelines.

### Tunicamycin injection and RNA extraction

We administered Tunicamycin (TM) or DMSO (control) (Sigma) with an intraperitoneal injection. TM was dissolved in DMSO (Sigma) to achieve a 2.5×10^-4^ mg/uL concentration. To induce ER stress in the mouse model, we injected mice with a final concentration of 1mg of TM per 1kg mouse weight (same concentration for DMSO control mice) [57, 58]. After injection, mice were allowed to recover for 8 hr. Mice were then euthanized and organs were harvested and stored at −80°C. RNA was isolated by Trizol (ambion) and Direct-zol RNA MiniPrep Kit (Zymo Research) RNA column extraction protocol.

### Illumina mRNA sequencing and alignment

mRNA sequencing was performed on 18 samples (3 genotypes X 3 replicates X 2 treatments) for liver and 18 samples for kidney, for a total of 36 samples. Samples were prepared and sequenced by the Huntsman Cancer Institute High-Throughput Genomics Core. Library prep was performed using Illumina TruSeq Stranded Total RNA Library Prep Ribo-Zero Gold. The 36 samples were then sequenced on the NovaSeq 2 x 50 bp Sequencing, for a total of approximately 25 million paired reads per sample. Fastq files were trimmed by using seqtk software. Parental RNA-seq reads were aligned to strain-specific reference genomes using Bowtie2 software [59]. Genomes were obtained from Ensembl (http://ftp.ensembl.org/pub/release-103/fasta/). F1 reads were aligned to masked genomes using STAR software, to allow for a more variant aware alignment [60]. Masked reference genomes were created using bedtools [61]. Within the B6 reference genome, known CAST variants are replaced with ambiguous N nucleotides. Alignment files were sorted and converted using samtools [62].

### Quantification of expression levels

The Deseq2 default normalization method was used to normalize counts [63]. For each genotype, condition, and tissue type, principal component analysis was used to identify outlying samples. For a given tissue, within a genotype, we required the TM samples to be clustered together and the control samples to be clustered together. If a given replicate was not with the appropriate cluster, it was removed from the analysis. At least two replicates remained after removing outliers for each combination of tissue, genotype, and condition. Remaining samples were reanalyzed using Deseq2. A gene was considered “expressed” if the Deseq2 value of base mean was ≥5.

### Effect of genetic background and tissue type on total expression levels

For each of the three genotypes, a single median expression value was calculated for each gene for control conditions, resulting in one control value per genotype. For each genotype, each replicate had a single TM value for each gene. This TM value was used to calculated TM-induced change in expression by taking the log2 ratio of the TM value over the median control value calculated earlier (log_2_(TM/control)).

This results in 2-3 values for each gene per genotype in liver and kidney. To test for a significant genotype effect on these values for each gene, we performed an ANOVA test. This was done for genes both upregulated and downregulated in response to ER stress in both liver and kidney, resulting in four different categories with genes that displayed strain-dependent expression.

### Allele-specific expression quantification in the F1 mouse

Allele-specific expression was quantified using GATK ASEReadcounter [64]. SNP information was obtained from the Sanger Mouse Genomes Project (https://www.sanger.ac.uk/data/mouse-genomes-project/). Counts for all replicates were combined to increase coverage and reduce variability. To increase the reliability of counts, we only included genes in our analyses that had at least 2 informative SNPs between B6 and CAST and at least 20 counts in at least one of the conditions. This resulted in 5,669 genes liver and 7,764 genes in kidney. Genes were considered to have a significant change in ASE post-ER stress if the ratio of the expression of the two alleles (B6 and CAST) under control conditions was significantly different than the ratio of expression under TM conditions determined by the Fisher’s exact test followed by a 5% FDR correction.

### Determination of regulatory mechanism

To determine whether the variable expression of a gene was due to *cis*-regulatory mechanism, *trans*-regulatory mechanism, or some other mechanism, we used a method that was previously described in McManus et al., 2010 and Chow et al., 2015 [8, 29].

This pipeline was run on counts for genes in each of the parental strains (B6 and CAST) as well as each of the alleles in the F1 mouse (B6 allele and CAST allele) for both control and TM conditions in liver and kidney. First, to determine if expression was significantly different between the two parental counts for a given gene, we performed a binomial exact test followed by a 0.1% FDR correction. Next, to determine if expression was significantly different between the two parental alleles in the F1, we performed a binomial exact test followed by a 0.1% FDR correction. Finally, to compare the ratio of parental expression to the ratio of allelic expression in the F1, we performed a Fisher’s exact test, followed by a 0.1% FDR correction. This results in two corrected binomial exact test p-values, and one corrected Fisher’s exact test p-value for each gene in the F1 for the control liver, control kidney, TM liver, and TM kidney. Based on these values, we categorized genes into: *cis-, trans-, cis-+trans-, cis-xtrans*-, compensatory, conserved, and ambiguous based on the guidelines listed below (S7 Tables).

#### cis-

Parental strains are different, alleles in F1 are different, but the ratio of parental expression compared to allelic expression is not different.

#### trans-

Parental strains are different, but the alleles in the F1 are not different, making the ratio of parental expression different than the ratio of allelic expression.

Next, for genes that displayed both *cis*- and *trans*-regulatory differences, they were divided into three categories: *cis-+trans-, cis-xtrans*-, and compensatory. Genes with *cis-+trans*-indicates both *cis*- and *trans*-mechanisms that favor the same allele. *cis-xtrans*-indicates both *cis*- and *trans*-mechanisms that favor opposite alleles. Finally, compensatory indicates evidence of both *cis*- and *trans*-mechanisms, but have no significant expression differences in the parental strains.

#### cis-+trans-

Parental strains are different, and parental alleles in the F1 are different. However, the ratio of parental expression and ratio of allelic expression is also different. The log of the ratio of parental expression was performed. A positive number indicates the B6 parental strain has a higher expression. Negative indicates the CAST parental strain has a higher expression level. Similarly, the log of the ratio of allelic expression was performed. Positive indicates the B6 allele has a higher expression level while negative indicates the CAST allele has a higher expression level. Genes where the higher expressed parental strain and higher expressed F1 allele matched, they were considered *cis-+trans*-(Ex: B6 parental strain & B6 allele had higher expression in each respective ratio). This indicates that both *cis- and trans*-mechanisms favor the same allele.

#### cis-xtrans-

Same criteria as *cis-+trans*-, except for the log of the parental expression and log of the allelic expression have opposite signs. This indicates that the *cis*- and *trans*-mechanisms favor expression of opposite alleles.

#### Compensatory

Parental strains are not different, but the alleles in the F1 are different, resulting in a difference in ratio of parental strains and ratio of alleles. This reveals that there is no significant difference in expression between the two parental strains despite evidence of both *cis*- and *trans*-regulatory mechanisms.

#### Conserved

There is no significant differences between parental strains, alleles, or ratios of both.

#### Ambiguous

Genes that do not fall into any of the previous categories.

Only genes that displayed *cis*- or *trans*-mechanisms are talked about and analyzed in the text.

### Enrichment analyses

All Gene Ontology analysis was performed with DAVID [65]. Transcription factor binding site enrichments were identified by using oPOSSUM [66]. We used the mouse single site analysis (SSA) tool with a cutoff of 2000 base pairs up- and downstream of the transcription start site.

## Acknowledgments

We thank Dr. Hans Dalton, Dr. Kevin Hope, Dr. Dana Talsness, and Maddie Haller for providing edits and suggestions for the manuscript. We thank Dr. Kwangbom Choi and Stephanie Kravitz for advice on ASE analysis. We thank Dr. Nathan Krah, Dr. Paul Bonthuis, Katie Owings, and Karen Peralta for assistance on mouse handling and dissections. We thank Dr. Charles Murtaugh for providing equipment for RNA extraction. We thank Dr. Rebecca Palu for technical and experimental assistance.

## Supporting information

**S1 Fig. Confirmation of ER stress induction.** *Xbp1* transcript is spliced by IRE1 when misfolded proteins are present in the ER. Spliced *Xbp1* transcript is a marker of ER stress. RT-PCR for spliced and unspliced *Xbp1* transcript in both control and TM mice for both liver and kidney (A). Upper band represents unspliced *Xbp1* at 183 bp and lower band represents spliced *Xbp1* at 157 bp. *Bip* is upregulated under ER stress conditions and is another hallmark of the UPR. *BiP* levels were measured by qRT-PCR in liver and kidney from control and TM injected mice (B).

**S2 Fig. Correlation of expression of common upregulated ER stress genes in each genotype.** Correlation of log_2_(TM/control) of the 566 common upregulated ER stress genes between liver and kidney in B6 (A), CAST (B), and F1 (C). Red line is regression line between the two tissues.

**S3 Fig. Gene expression correlation of common downregulated ER stress genes in each genotype.** Correlation of log_2_(TM/control) of the 106 common downregulated ER stress genes between liver and kidney in B6 (A), CAST (B), and F1 (C). Red line is regression line between the two tissues.

**S4 Fig. Genes that show the same *cis*- or *trans*-regulatory mechanism under both conditions are strongly correlated in their magnitude.** Data for liver (A) and kidney (B). ER stress does not affect the regulation of these genes (Liver: *cis*-: 223, *trans*-: 104; Kidney: *cis*-: 351, *trans*-: 64).

**S5 Fig. Genes that show the same *cis*- or *trans*-regulatory mechanism in both tissues are slightly correlated in their magnitude.** *cis*-mechanisms observed in tissues under control (A) and TM (B) conditions. *trans*-mechanisms in both tissues under control (C) and TM (D) conditions.

**S6 Fig. No genotype bias in genes with significant change in ASE.** The proportion of genes with an ER stress-induced change in ASE that show an increase of the B6 or CAST allele in liver (A) or kidney (B). No genotype bias in genes that display significant change in ASE and significant change in RNA transcript levels in liver (C) or kidney (D). For genes downregulated in kidney, there is a trend towards the CAST allele contributing more towards these ASE effects. However, due to the low number of genes in this category, this bias is not significant (p=0.136).

**S7 Fig. The majority of genes showing ER stress-induced ASE are tissue-specific.** The proportion of genes that display a significant change in ASE post-ER stress that in liver, kidney, or both.

**S1 Table. ER stress induced upregulated genes.**

**S2 Table. ER stress induced downregulated genes.**

**S3 Table. GO enrichment analysis of ER-stress induced genes.**

**S4 Table. Tissue-specific GO enrichment.**

**S5 Table. ER stress-induced genes with strain effect.**

**S6 Table. GO enrichment analysis of strain-dependent genes.**

**S7 Table. Counts for each regulatory mechanism.**

**S8 Table. B6xCAST F1 liver.**

**S9 Table. B6xCAST F1 kidney.**

**S10 Table. Allele specific expression.**

**S11 Table. Transcription factor binding site enrichment analysis.**

