## Supplemental Figures for "The dynamic effect of regulatory genetic variation on the *in vivo* ER stress transcriptional response"

**S1 Fig. Confirmation of ER stress induction.**

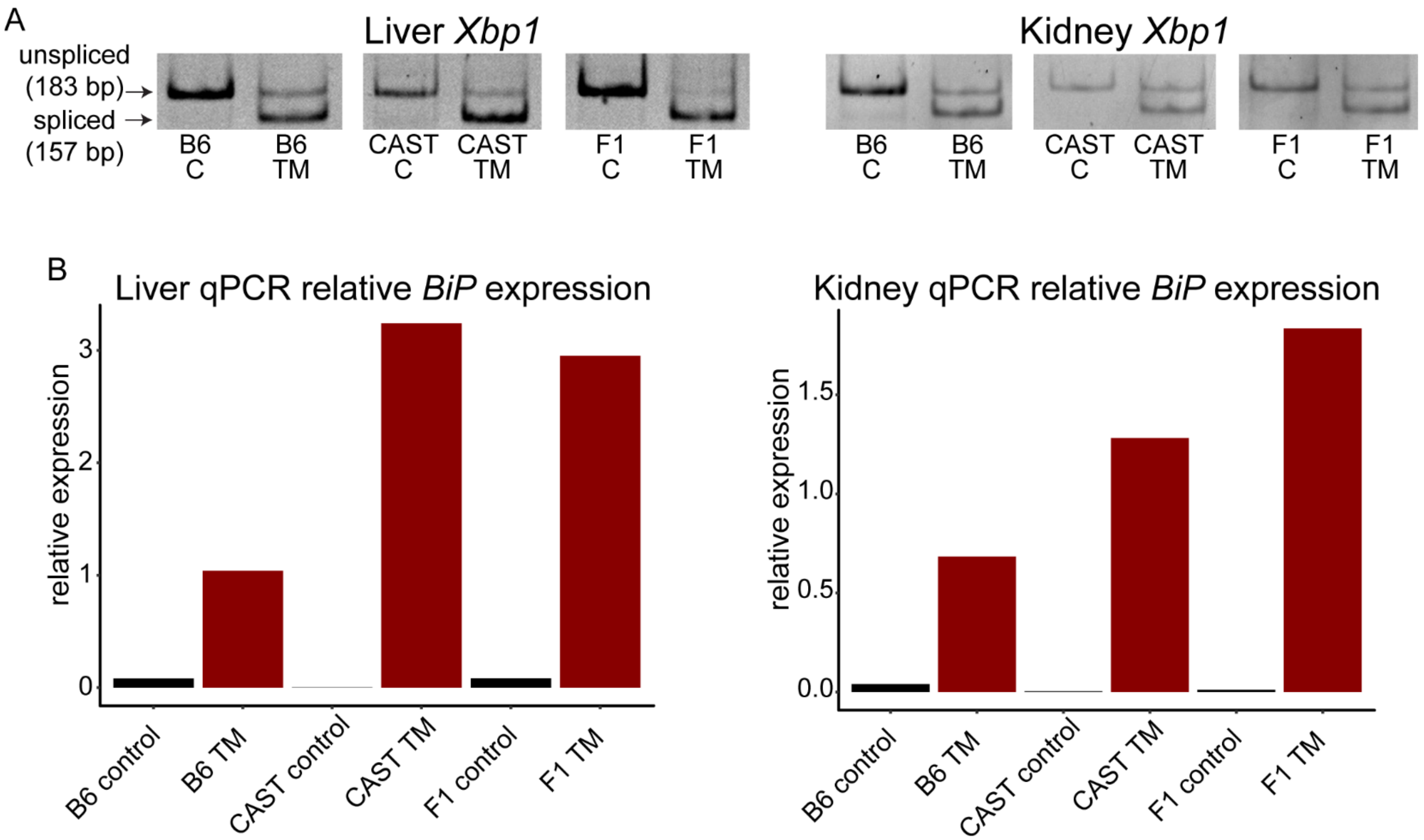

**S2 Fig. Correlation of expression of common upregulated ER stress genes in each genotype.**

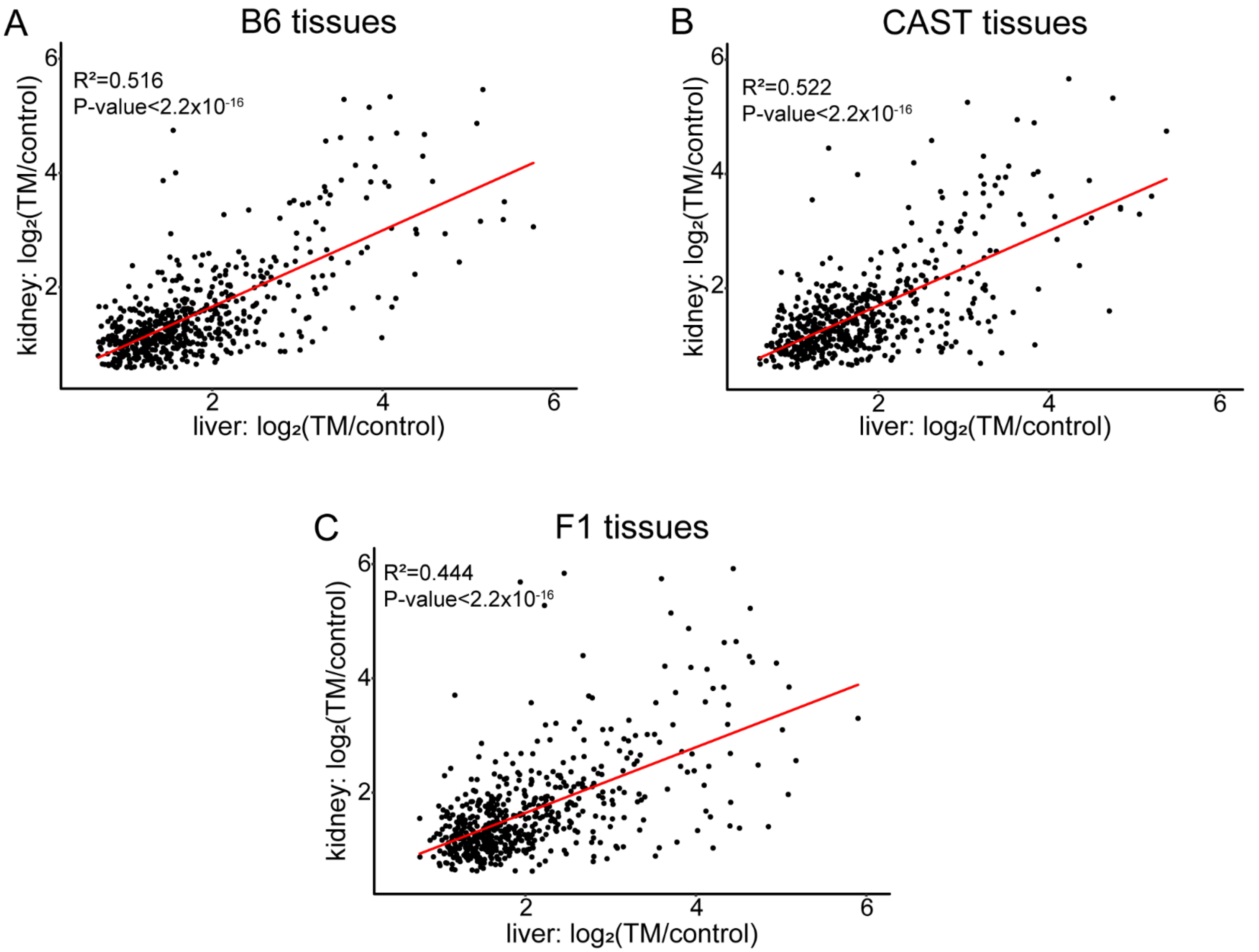

**S3 Fig. Gene expression correlation of common downregulated ER stress genes in each genotype.**

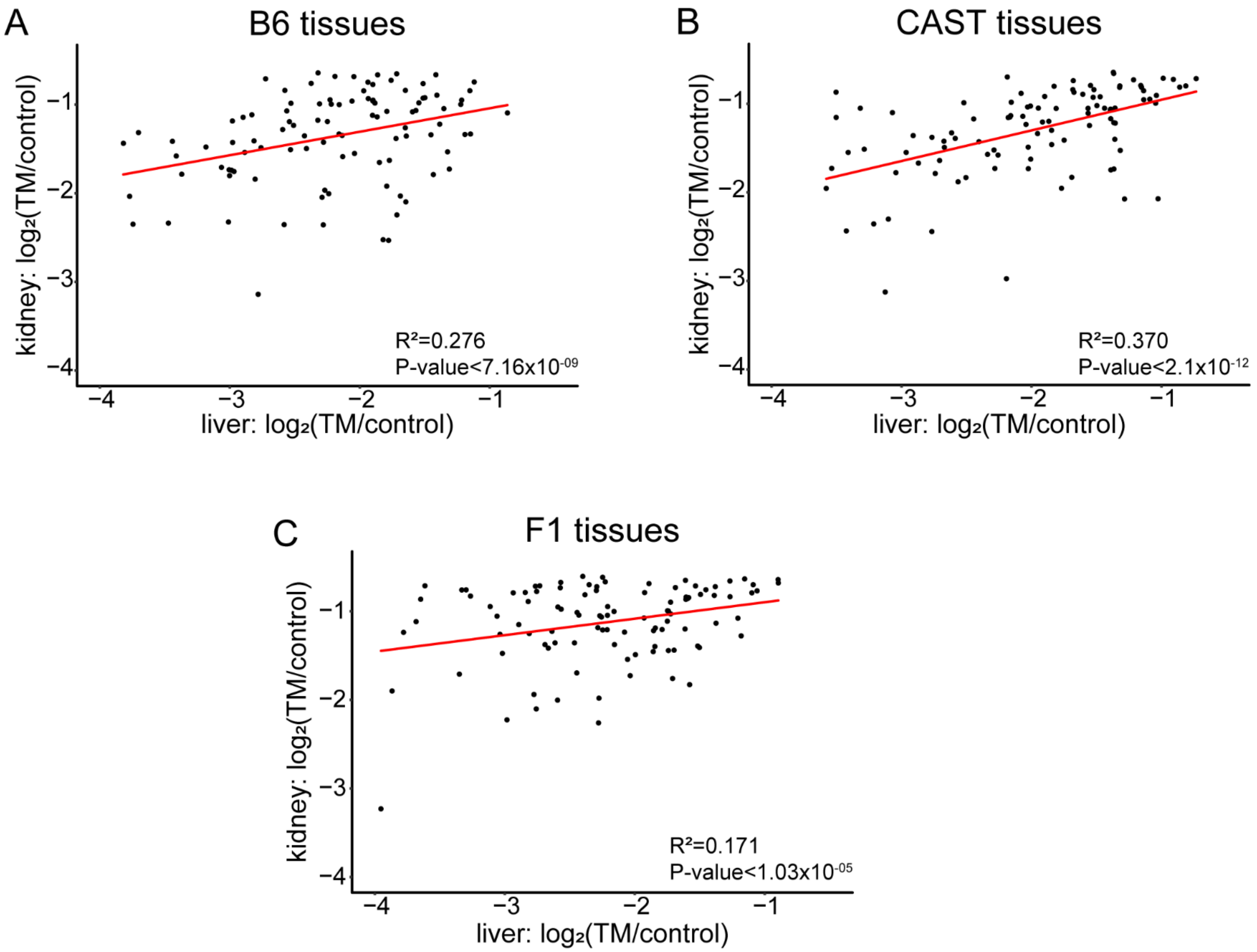

**S4 Fig. Genes that show the same *cis-* or *trans-* regulatory mechanism under both conditions are strongly correlated in their magnitude.**

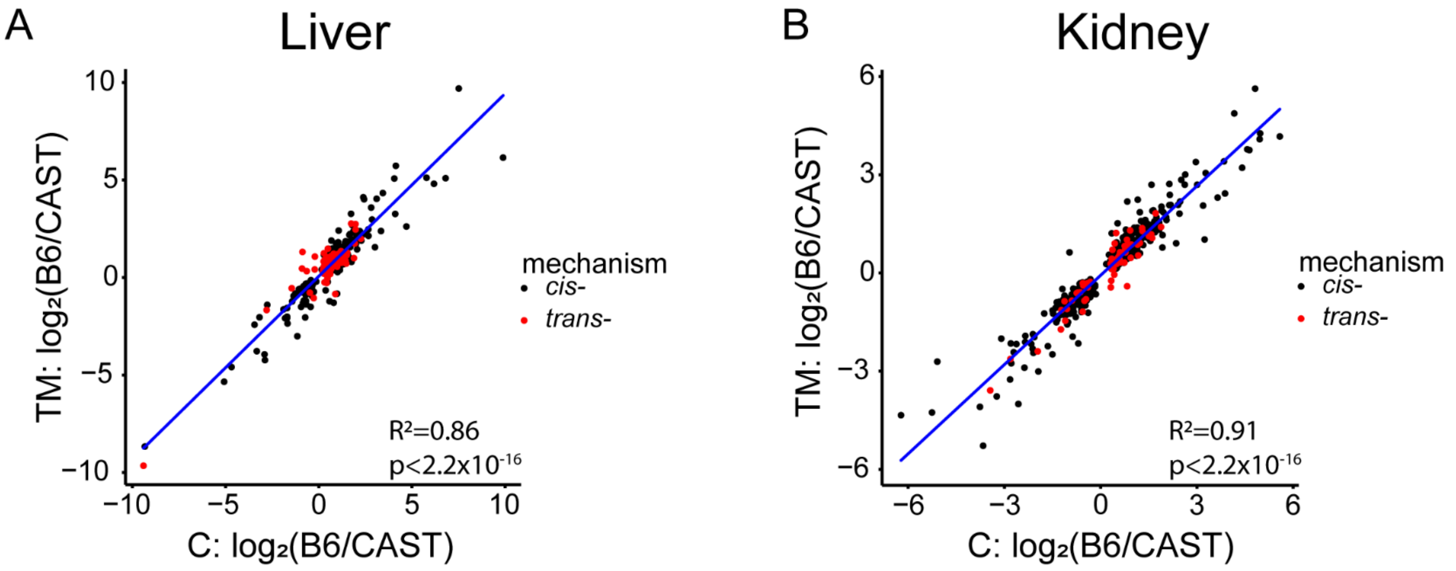

**S5 Fig. Genes that show the same *cis-* or *trans-* regulatory mechanism in both tissues are slightly correlated in their magnitude.**

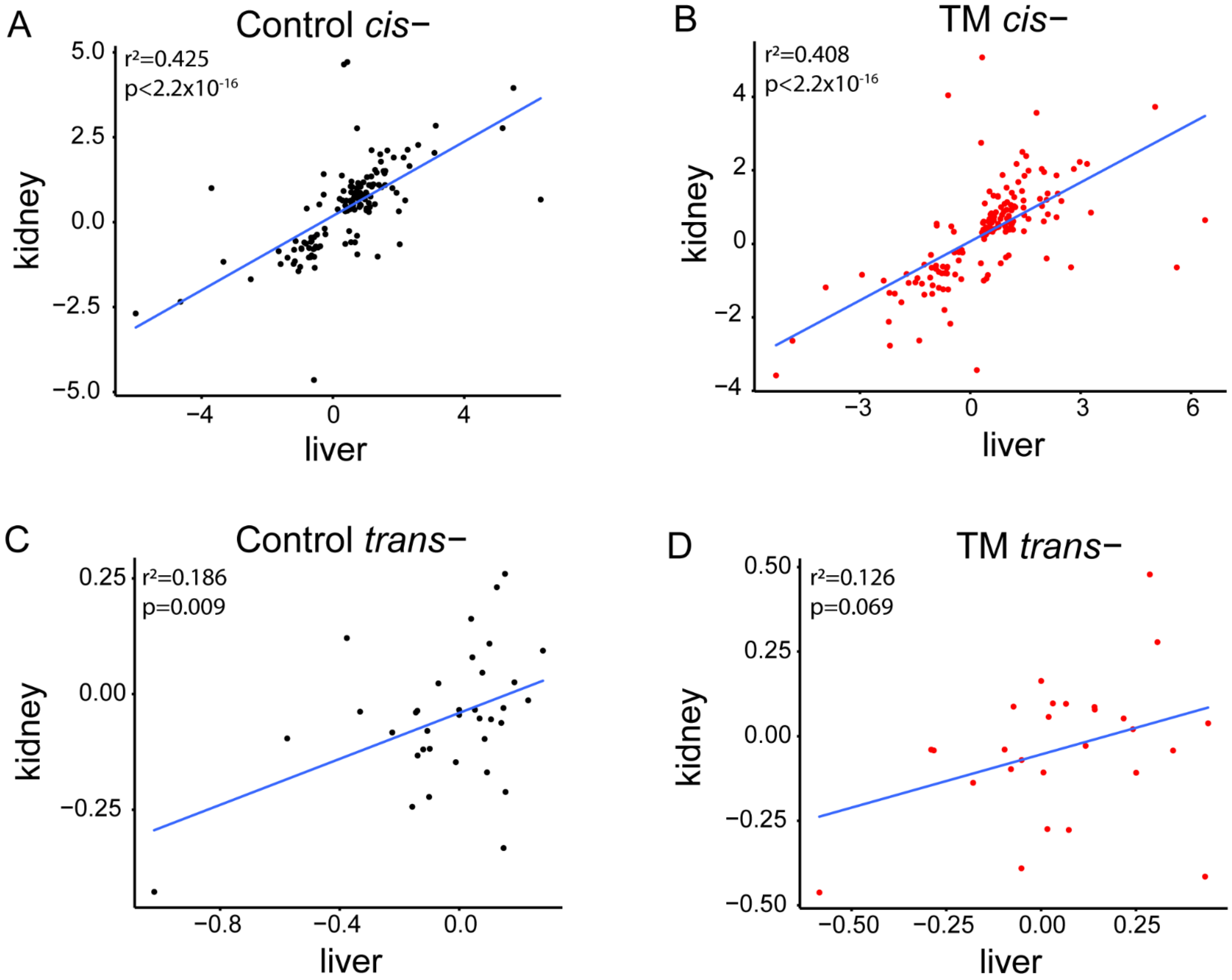

**S6 Fig. No genotype bias in genes with significant change in ASE.**

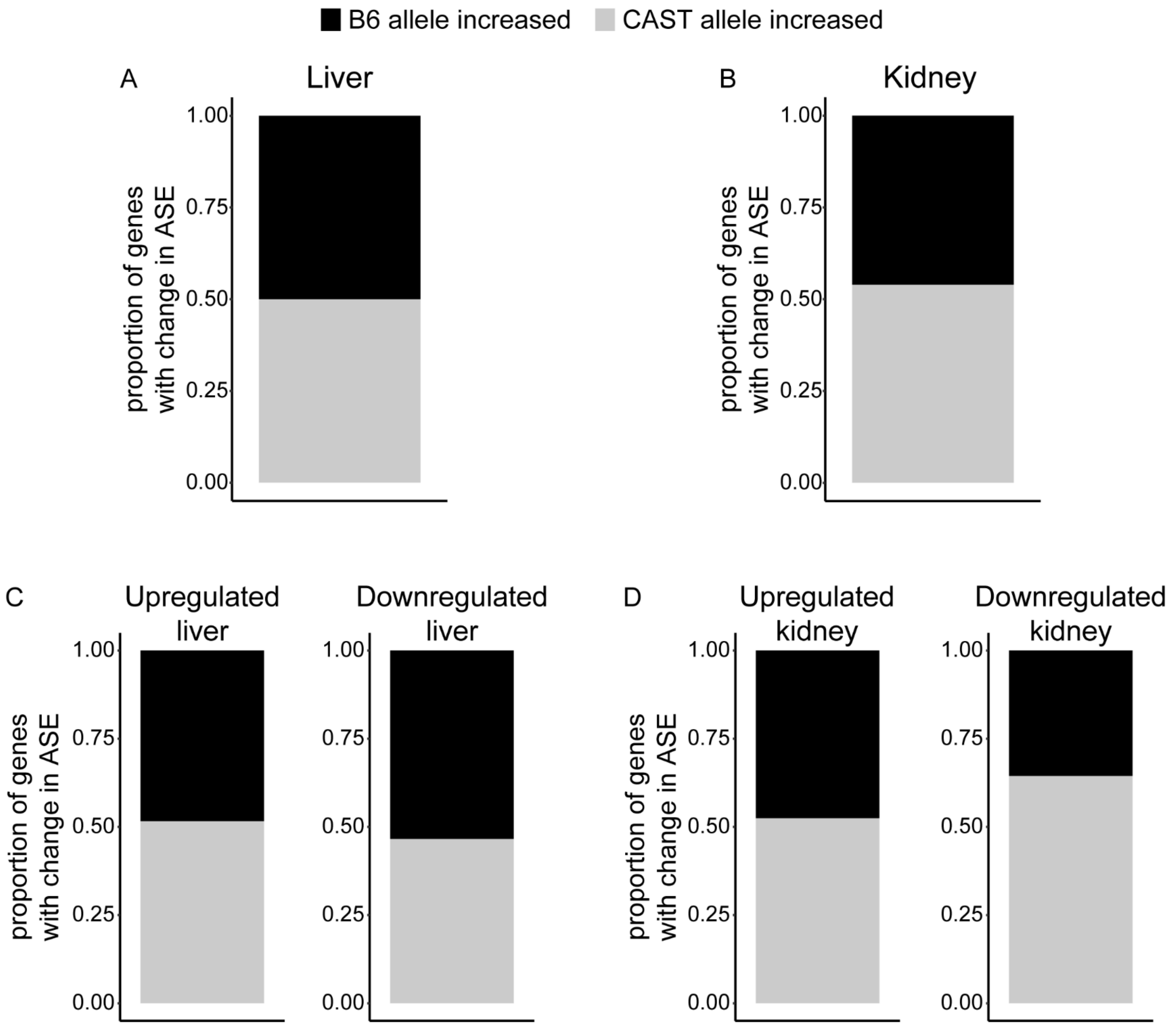

**S7 Fig. The majority of genes showing ER stress-induced ASE are tissue-specific.**

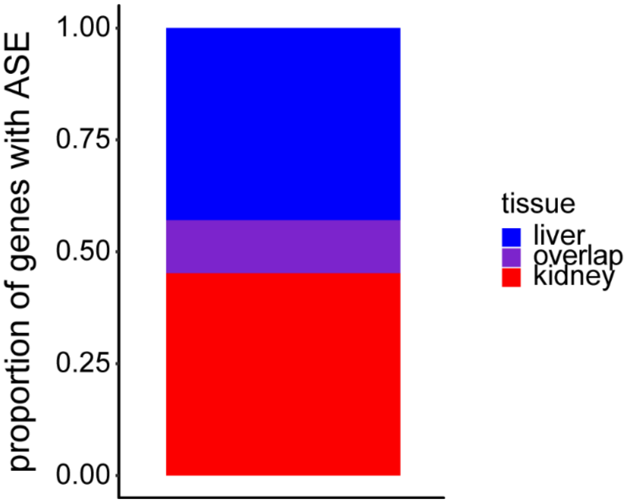
